# Defining a Muscle Stem Cell matrisome signature: from transcriptome data to extracellular matrix niche topology

**DOI:** 10.1101/2024.07.10.602878

**Authors:** Emilie Guillon, Bacar Hisoilat, Takako Sasaki, Philippos Mourikis, Florence Ruggiero

## Abstract

Although intensively investigated, the regulation of skeletal muscle stem cells (MuSCs) by their niche remains an open question. The extracellular matrix (ECM) components of the niche represent a dynamic microenvironment that undoubtedly participates in MuSCs behavior. We used bioinformatics analysis of transcriptomic data to define the matrisome profile of skeletal muscle resident cells, comprising genes encoding ECM and ECM-associated proteins. We identified quiescent MuSCs as key ECM producers of the niche, notably through the expression of specific basement membrane genes as *Col19a1* and *Lama3* and regulators of ECM assembly, *Thsd4 and Aebp1*. Unexpectedly, quiescent MuSCs also expressed matrisome neurogenesis-related genes. Immunofluorescence staining of selected ECM components showed their organization in isolated murine myofiber bundles. Upon activation, MuSCs strikingly downregulated the niche-related ECM genes and instead expressed genes involved in basement membrane disruption and matrisome genes linked to cell motility. This study identified distinct matrisome signatures of quiescent and activated MuSCs that are consistent with their function in homeostasis and repair of damaged skeletal muscle.

## INTRODUCTION

The skeletal muscles are responsible for almost all body movements and represent 40% of total human body weight. Besides the myofibers that are the major cell components of skeletal muscles, several other cell types including tissue-specific cells are present such as the fibro-adipogenic progenitors (FAPs), adipocytes, macrophages, and importantly the muscle stem cells (MuSCs) also known as satellite cells (Relaix et al., 2021). Due to the coordinated activity of these cells, skeletal muscles show remarkable capability of regeneration. Cell depletion experiments have shown that MuSCs are the indispensable cells for tissue repair (Sambasivan et al., 2011, Lepper et al, 2011). In the adult muscle, MuSCs are marked by the paired homeobox transcription factor Pax7 and are mitotically quiescent. In response to injury, they get activated, downregulate Pax7 and enter the cell cycle. Proliferating myoblasts either self-renew to maintain the MuSC pool or differentiate and form new myofibers.

Adult stem cells reside in a specific microenvironment known as the niche whose main function is to maintain quiescence and control cell fate. Specific changes in the composition and/or integrity of the niche can result in the activation and proliferation of stem cells. In resting adult skeletal muscle MuSCs are wedged between the basement membrane (BM) of the underneath myofiber and its sarcolemma. The BM constitutes the direct MuSC environment and is thus considered as a critical part of the MuSC niche that can regulate MuSC fundamental properties including proliferation, self-renewal, differentiation and migration by providing specific signals to the cells (Rayagiri et al., 2018). However, the extracellular matrix (ECM) composition of the niche and the niche-specific role of ECM on MuSC behavior remain poorly documented.

Recent studies have highlighted the role of collagens in MuSC behavior. Urciuolo and collaborators showed that collagen VI regulates MuSC renewal in mice. Skeletal muscle regeneration after injury is impaired in mice lacking *Col6a1* gene and the self-renewal capacity of MuSCs is considerably reduced in absence of this collagen (Urciuolo et al., 2013). Collagen V plays instead an important role in the maintenance of MuSC quiescence in a cell-autonomous fashion through a Notch-ColV-Calcitonin receptor signaling cascade (Baghdadi et al., 2018). Interestingly, our laboratory showed that these two collagens interacted together and were part of a same ECM network in the skin (Bonod-Bidaud et al, 2012; Symoens et al., 2010), raising the possibility of a coordinated action of COLV and COLVI on MuSC regulation. The matrisome was defined by Naba and co-workers as the ensemble of genes encoding secreted proteins that are found in the extracellular space (Naba et al., 2012). Based on proteomics methods coupled with bioinformatics, they were the first to characterize the protein composition of tissue extracellular proteins in humans and mice. They described 1110 and 1027 genes for the mouse and human matrisomes, respectively, which were subdivided into two main categories: the core matrisome comprising EMC structural proteins (collagens, proteoglycans and glycoproteins), and the matrisome-associated that includes all other secreted proteins and is sub-divided into EMC regulators, secrete factors and EMC-affiliated proteins.

The composition and topology of the MuSC niche, as well as the cell contribution in building-up the niche are far from being fully elucidated. To fill this gap, we used transcriptomics data (bulk and single-nucleus RNAseq) of skeletal muscle cells to characterize the quiescent and activated MuSC matrisomes. Bioinformatics analyses revealed specific profiles of quiescent MuSCs with core-matrisome specificities and an unexpected enrichment of “neurogenesis-related” genes that are dysregulated upon activation. Light-sheet microscopy analysis of intact myofiber bundles with antibodies to selected ECM structural proteins showed particular topology of ECM components of the niche and identified new markers of MuSCs.

## RESULTS

### Quiescent MuSCs are a major source of matrisome genes including core-matrisome genes

To characterize the matrisome of quiescent MuSC (qMuSC), we analyzed transcriptomic data of *in situ* fixed cells, which we have previously reported (Machado et al., 2017). In that study, intact muscles of *Tg:Pax7*-nGFP mice were fixed with paraformaldehyde before enzymatic treatment to avoid MuSC activation during the cell isolation procedure. We extracted the matrisome gene set expressed by qMuSCs using the online mouse matrisome databank (Naba et al., 2012; http://matrisomeproject.mit.edu/).

Among the 1110 matrisome genes present in the mouse genome, qMuSC expressed 242 genes representing no less than a fifth of the mouse matrisome. Further analysis of the relative proportions of each matrisome category showed that qMuSCs produce a high percentage of core matrisome genes (37%) compared to other sub-categories (Figure 1A). This result was confirmed when core matrisome and matrisome-associated genes in qMuSCs are calculated as percentages of the total number of genes in the mouse matrisome. qMuSCs express 32% of the core matrisome potentially available in mouse genome compared to 18% of the matrisome-associated genes (Figure 1B). A detailed analysis of the core matrisome sub-categories (including collagens, glycoproteins and proteoglycans) revealed that qMuSCs surprisingly expressed a high number of collagen genes (19 of the 44 collagen genes in the mouse genome), which represents 43% of this sub-category compared to 32% for glycoproteins and 19% for proteoglycans (Figure 1B). This observation is strengthened by the fact that 9 genes encoding collagen α-chains are found in the 50 top matrisome genes expressed by qMuSCs (Figure 1C, Table S1). Notably, among these 9 genes are collagen V and collagen VI genes, which is consistent with the known role of these collagens in MuSCs (Baghdadi et al., 2018, Urciuolo et al., 2013). Collagen III, IV, XV and XIX genes not previously described as being expressed in quiescent or activated MuSCs were also identified. At the structural level, these collagens are components of the basement membrane (BM), like the basement membrane associated zone (BMZ) collagens IV, XV and XIX, or of the interstitial ECM, like fibrillar collagens V and III. Collagen V was initially identified at the interface of the BM and interstitial ECM in different tissues (Ricard-Blum and Ruggiero, 2005; Bonod-Bidaud et al., 2012). Gene ontology (GO) enrichment analysis on qMuSC matrisome also identified structural ECM terms as over-represented, including BM genes (Figure 2A, B). qMuSC matrisome was indeed significantly enriched in the gene sets “Extracellular matrix“, “Collagen containing matrix” and “Basement membrane” (Figure 2A, green box). Gene sets associated with cell-ECM interactions such as “cell adhesion”, “biological adhesion”, “integrin binding”, “cell adhesion molecule binding”, “extracellular matrix structural constituent” and “extracellular matrix structural constituent conferring tensile strength” were likewise enriched (Figure 2C, green box).

**Figure 1.**
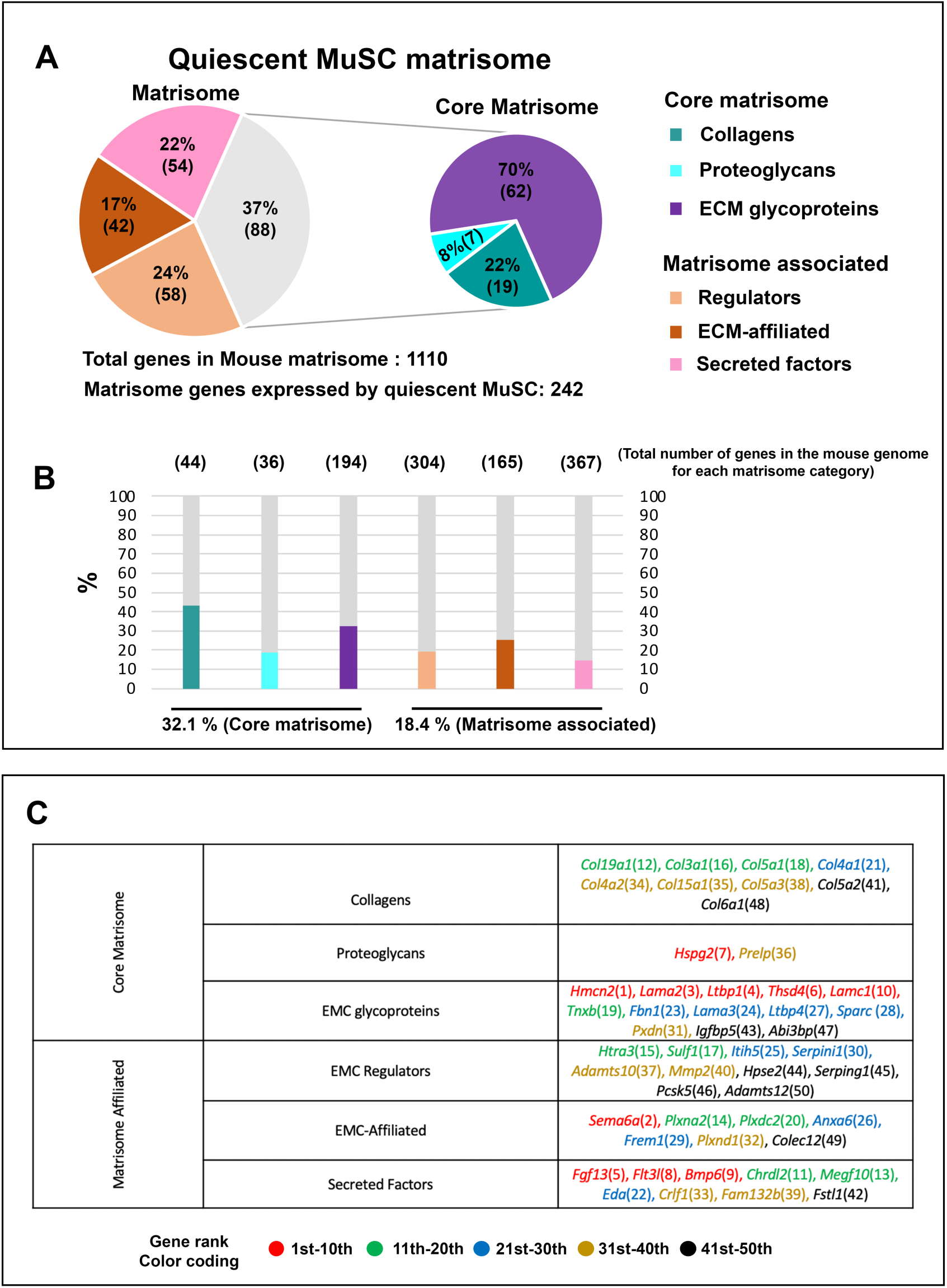
Characterization of quiescent MuSC Matrisome. (A) Pie chart represents the number genes expressed genes for each matrisome category and the relative proportion of each category in the total matrisome or core matrisome expressed by qMuSC. (B) Proportion of genes expressed by quiescent MuSC relative to total number of genes available in the mouse genome for each matrisome category. Total number of gene in the mouse genome for the corresponding category is indicated in brackets above the histogram. (C)Top 50 of matrisome genes expressed by qMuSCs. Ranks are indicated into brackets.

**Figure 2.**
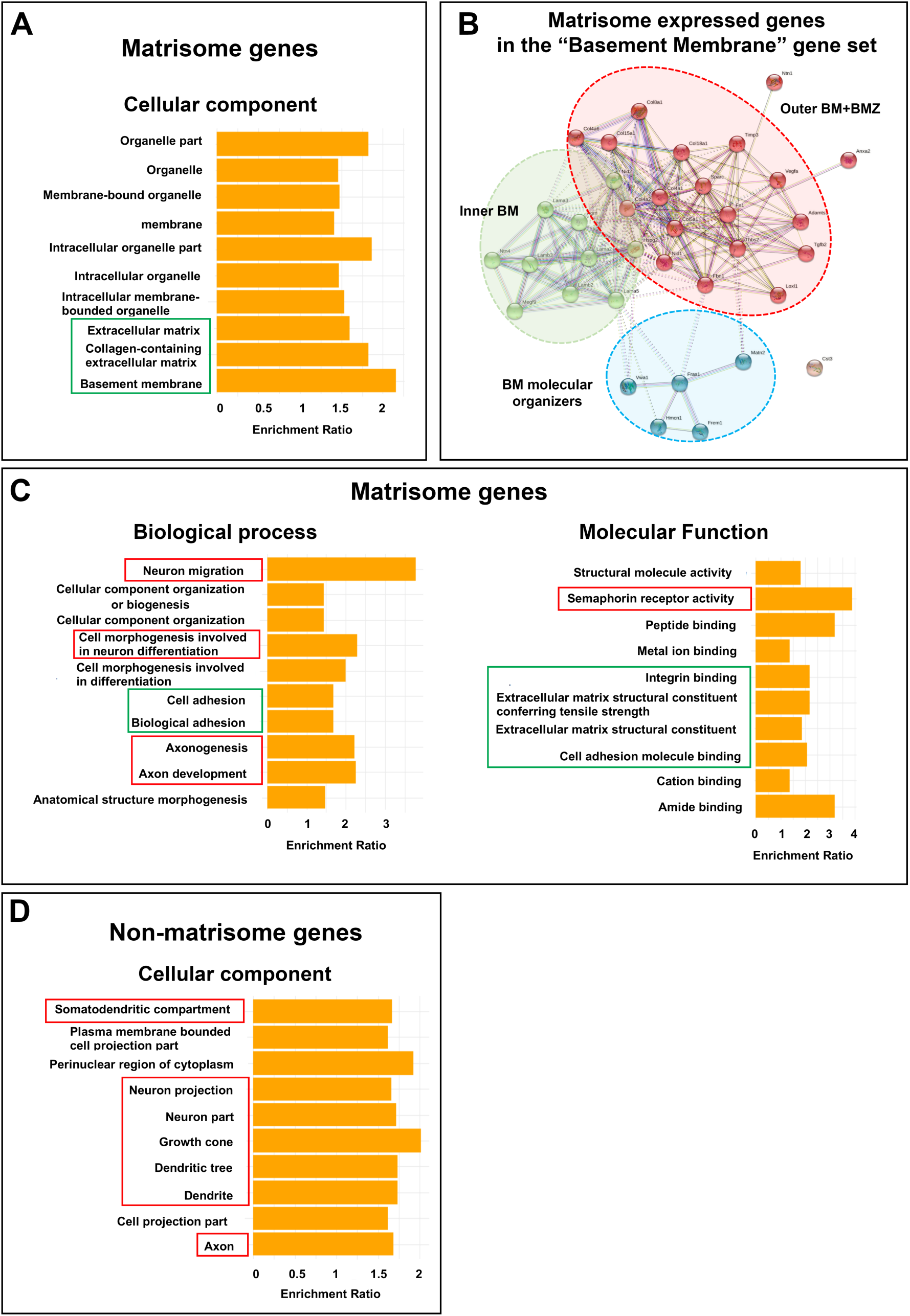
Quiescent MuSCs express basement membrane and neurogenesis-related tool-kit genes. (A,C,D) Gene ontology enrichment analysis of matrisome or non-matrisome expressed genes by qMuSCs; Cellular component (A,D), Biological process (C) or Molecular function (C) domains were analyzed. Only Top 10 enriched GO terms are shown. GO terms linked to “Extracellular matrix” or “Neurogenesis” features are highlighted in Green or red respectively. (B) Protein-protein interaction network analysis of the genes mapped on the “basement membrane” gene set in A was done with STRING database. The 35 expressed matrisome genes were input into STRING database and created a network with an enrichment p-value <10^-16^. Clusters were generated using a MCL parameter of 2 (one color represents one cluster).

The MuSCs lie directly underneath the myofiber basement membrane that is part of its niche composition. The enrichment in the GO term “basement membrane” that shows the highest score was thus of particular interest. In support to this finding, several BM components were found in the top 50 expressed genes (*Lama2, Lama3, Lamc1, Col4a1, Col4a2, Hspg2*) (Figure 1C and Table S1) corroborating the fact that qMuSCs actively contribute to the BM maintenance. Among the 71 genes of this gene set, qMuSC was shown to express 35 BM-related genes. Interestingly, STRING interaction network analysis on these 35 genes revealed 2 major distinct hubs consisting in genes involved in the formation of the inner BM and the outer BM and BMZ, respectively (Figure 2B). The “Inner BM” hub contains laminin genes and those involved in the formation of the laminin network. The “Outer BM and BMZ” hub comprised several collagen genes including collagen IV α-chain genes and BMZ collagen genes such as *col15a1* and *col18a1*, but also regulators of BM structure as *Sparc* that is known to modulate collagen IV levels in the BM (Morrissey et al., 2016). Of interest, *Nid 1, Nid 2* and *Hspg2* were found at the interface of the 2 hubs, consistent with their linker function in the BM as nidogen and perlecan interact with both laminin and collagen IV networks. Lastly, a small but important third hub appeared containing several gene coding for proteins involved in the BM organization and BM/BM linkage such as *Frem1*, *Fras1* and *Hemicentin-1* (*Hcmn1*) (Pavlakis et al., 2011; Keeley and Sherwood, 2019).

In addition, although absent from the current gene list of the GO term “Basement membrane”, the BMZ components, *Col19a1* (collagen XIX) and *Hcmn2* (Hemicentin-2) (Calvo et al., 2020, Keeley and Sherwood, 2019) were both found to be highly expressed in qMuSCs (Figure 1C and Table S1). *Hcmn2* was the most highly expressed matrisome gene of qMuSCs displaying a read count 20 times higher than the last gene from the top 50 (Table S1). In conclusion, the bioinformatics analysis of the qMuSCs RNA-seq transcriptome data revealed qMuSCs as potential architects of their own niche, contributing to the building-up and remodeling of the BMZ and beyond as they also express interstitial ECM genes.

### Quiescent MuSCs express matrisome genes classically involved in extracellular regulation of neurogenesis

Besides this structural aspect, our analysis revealed another unexpected feature of the qMuSC matrisome as GO terms related to the regulation neurogenetic processes were found to be highly enriched. As such, Semaphorin 6A and its receptors Plexin A2 and D1 were found in the qMUSC top 50 expressed genes (Figure 1A). Semaphorins are a large family of secreted, transmembrane, or GPI-anchored proteins that are known to control a wide variety of developmental processes in the nervous system such as axon guidance, cell migration, dendrite morphology, and synaptogenesis through their receptors neuropilins and plexins (Verhagen and Pasterkamp, 2020).

An in-depth analysis showed that, although at a lower level, qMuSCs expressed six other semaphorin genes and 8 out of 9 plexin genes contained in the mouse genome (Figure 2C, right panel). In addition to “Semaphorin receptor activity genes” 4 other GO terms related to neurogenetic processes were found to be enriched: “neuron migration” that shows the highest enrichment score, followed by “cell morphogenesis involved in neuron differentiation”, “axonogenesis” and “axon development” (Figure 2C, left panel). Among the genes contained in these GO terms, several genes encoding proteins with well documented functions in neurogenesis such as *Agrin* involved in the neuromuscular junction formation, *Slit 2* and *Slit 3* that are key axonal guidance cues, and *Tenascin C* and *Tenascin R* involved in neurogenesis (Iversen et al., 2020; Tucic et al., 2021).

Next, the question of whether this tendency to express genes associated with neurogenesis is a matrisome specificity was addressed. To test this, a GO analysis of the non-matrisome genes expressed by qMuSCs (all genes except the matrisome ones) was conducted. Contrary to what observed for the qMuSC matrisome, non-matrisome genes expressed by qMuSC did not show enrichment in biological processes and molecular functions related to neurogenesis (data not shown). However, analysis of the cellular component sub-ontology category of non-matrisome genes (Figure 2D) revealed a striking enrichment in gene sets related to cellular structures specific to neural cells: 7 enriched gene sets were related to neuron cell projections (axonal or somatodendritic) or axonal growth cone and 2 other enriched gene sets were related to cell projection in general.

Taken together the data showed that qMuSCs produce an unconventional gene toolbox classically associated with shaping neuronal cells morphology. How the encoded proteins of these genes have adapted to the shape and function of qMuSCs remains an open and intriguing question.

### Quiescent MuSC display high morphological variability

Before focusing on ECM topology, we analyzed in more detail the morphology of MuSC in respect to their long projections, which is a hallmark of quiescence (Verma et al.,2018; Ma et al.,2022; Kann et al.,2022) (Figure S1). Recent studies have described the function and dynamics of these projections as well as imaging of whole mount muscles. We further explored those projections and sorted MuSCs in 6 different profiles based on their number, length and orientation (Figure S1A). When quantifying the proportion of MuSC displaying each profile, we found that the standard morphology with 2 long symmetric protrusions (a.k.a. quiescent projections) was not the most frequent one (19%). Instead, MuSCs with 1 long protrusion or with asymmetric protrusions were predominant (36% and 26% respectively) (Figure S1B). Morphologies with no protrusions, 1 short protrusion and atypical protrusions were less frequent (8%, 6% and 5% of MuSC cells, respectively).

The high morphological variation suggests that the projections of qMuSCs are more dynamic than previously thought. While intravital imaging strongly suggests that qMuSC are anchored to a fixed position, the extent to which the projection change length and shape remains ill-defined (Webster et al. 2016; Ma et al., 2022). Instead, it has been clearly demonstrated that the projections are retracted rapidly upon activation (Ma et al., 2022; Kann et al., 2022). Finally, we observed 8% of MuSCs without projection, likely corresponding to the small fraction of primed cells that are found in homeostasis (Snijders et al., 2015)

### MuSC niche is associated with fibrillar Collagen I rich areas but not with BM-specific features

Several of the ECM proteins that are robustly expressed by the qMuSCs are also found in the myomatrix (Csapo et al., 2020). However, the 3D network organization of these ECM components along the myofiber and particularly in the MuSC niche have not been systematically characterized. We speculated that a specific ECM structure and organization in the niche topology could favor MuSC quiescence.

To test this assumption and provide a comprehensive view of the 3D topology of ECM components of the qMuSC niche, we developed an imaging protocol based on mechanical dissociation and light-sheet imaging of whole muscle bundles. Co-staining of MuSC and ECM components that are commonly performed on transverse cryosections makes 3D reconstruction very challenging. Likewise, staining of single myofibers isolated from muscle using collagenase digestion may not reflect native collagen networks at interfaces. Indeed, mechanical dissociation has been recently shown to better preserve native MuSC morphology and BM network topology compared to collagenase-based isolation protocols classically used so far in the field (Schüler et al., 2021; current protocols). Here, fixed myofiber bundles were mechanically dissociated and co-immunostained for various ECM components and the cell surface marker M-cadherin to visualize MuSC morphology. Bundles were imaged with a light-sheet microscope and 3D reconstruction was performed from images taken inside the bundles to ensure analysis of intact interfaces (Figure 3A-D, schema in E, left panel).

**Figure 3.**
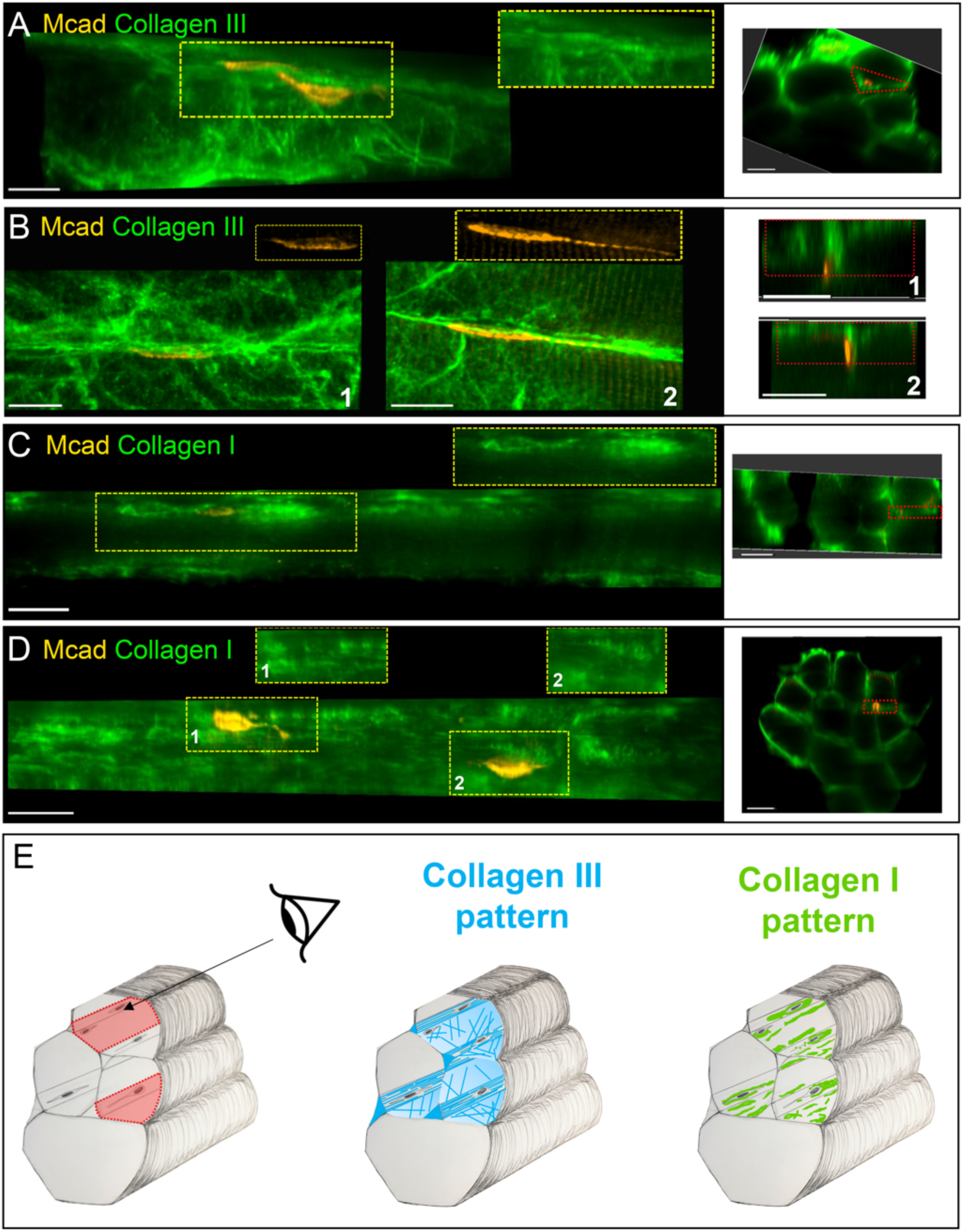
MuSCs co-locate with ECM zones enriched in Fibrillar Collagen I. (A-D) Co-immunostaining of mice muscle bundles (8-12 weeks old) with anti-M cadherin (Mcad) to label MuSCs in orange and Collagen III (A,B) or Collagen I (C,D) antibodies (green). Projected top views of 3D myofiber interfaces isolated as illustrated in the schematic in F (left panel). The corresponding transverse sections indicate the position of each interface in the whole muscle bundle (red rectangle). All images were acquired with a light-sheet microscope except in B which were acquired by classical confocal microscopy. Cropped areas in A,C,D show regions in the corresponding yellow frames without MuSCs to visualize matrix signal at the cell position. Cropped areas in B show the Mcad signal only to better appreciate MuSC morphology. Scale bars= 20 µm in A,15 µm in B and 30 µm in C,D. (E) 3D Schematic of a muscle bundle with MuSCs cells and representative patterns of collagen III (middle panel) or collagen I (right panel) at muscle fiber interfaces. In red on the left panel, are indicated myofiber interfaces as they are seen in A-D.

We analyzed collagen III topology. Collagen III organized into a complex filamentous network in which multi-directional and long filaments were clearly visible along the myofibers (Figure 3B, C and graphical representation in Figure 3A middle panel). MuSCs were observed in close contact with collagen III network suggesting that it could be a component of their niche. However, no obvious organization of collagen III at the MuSCs proximity was observed compared to the rest of the muscle bundle interfaces (Figure 3A-C). It is known that Collagen III and Collagen I can assemble to form mixed fibrils (Ricard-Blum and Ruggiero, 2005). In addition, it was shown that qMuSC express *col1a1* gene (Baghdadi et al., 2018) even though it was absent in our gene expression profile analysis as it was under the 1000 reads cut off (436 reads). Therefore, we decided to analyze Collagen I topology and showed that its deposition displays a very different pattern than Collagen III suggesting that at least some fibrils are not heterotypic Collagen I/III fibrils. Collagen I showed an unexpected heterogenous deposition along the myofiber interfaces forming a patchy pattern with noticeable enriched areas at the interfaces (Figure 3 C, D and graphical representation in Figure 3E right panel). Remarkably, patches rich in collagen I were observed at the location of MuSCs: 74.5% of cells (n =137 MuSC, pooled data from 8 muscle bundles from 3 muscles of 3 different mice) were surrounded (Figure 3C and 3D, cell 1) or at least bordered on one side (Figure 3D, cell 2) by a patch of collagen I (Figure 3D).

In contrary, the BM collagen IV (Figure S2A) and BMZ collagen XV (Figure S2 B, C) staining did not reveal any specific organization or enrichment at the MuSC niche. A strong enrichment was instead observed around blood vessels consistent with their known functions in angiogenesis (Figure S2A-C). Collagen VI immunoreactivity was observed along the myofiber interfaces but showed a high to low intensity gradient from the periphery to the center of the bundle myofiber interfaces (Figure S2D). No specific organization of collagen VI at the MuSCs niche was visible at the light-sheet microscope resolution. Unfortunately, in our experimental conditions, staining with commercially available anti-HCMN2 antibodies showed non-specific staining.

In conclusion, collagens I, III, IV, XV are abundantly present on the myofibers, surrounding the MuSC vicinity. While these ECM proteins in muscle may originate from various cellular sources, including interstitial mesenchymal cells, the qMuSC transcriptome indicates that qMuSC themselves express these proteins robustly. This suggests a potential specialized role for these proteins within the MuSC niche.

### Quiescent MuSC display a specific matrisome signature

The fact that most of the collagens we immunostained in this study seemed to be equally distributed along the myofiber without any particular enrichment or organization around MuSC raised the question whether and to what extent other neighboring cell types contributed to the ECM environment. Notably, FAPs that are in close proximity to MuSCs are described as major EMC producers in skeletal muscle (Figure S3A). Myofibers to which MuSCs are directly anchored are also producing ECM, mostly BMZ components. MuSCs are also very often in close proximity to blood vessels (white stars, Figure S2C, D) therefore endothelial cells could also be contributing to the MuSC niche.

To answer this question and potentially identify a specific matrisome gene expression signature associated to MuSCs, we conducted a comparative pairwise analysis of the matrisomes expressed by FAPs *vs* MuSCs, Myofibers *vs* MuSCs, and endothelial cells *vs* MuSCs. With this aim, we took advantage of single-nucleus RNA sequencing (snRNAseq) data from intact murine skeletal muscle that we have reported previously (Machado et al., 2021). This “intact muscle” dataset contains nuclei from 10 different cell populations. The gene expression profiles from FAPs, MuSCs, endothelial cells and the major myonuclei cluster referred to as generic Myonuclei (gMyonuclei) (as opposed to specialized nuclei located at neuromuscular and myotendinous junctions) were compared. Matrisomes were characterized for each nucleus population and the differential expression levels of core matrisome genes and matrisome-associated genes were plotted (Figure S3B, C). FAPs, MuSC, endothelial cell, and gMyonuclei were found to express 436, 357, 313, and 266 matrisome genes, respectively. Differential gene expression analysis of the structural components of the matrisome between FAP and MuSC showed that most of the core-matrisome genes expressed by MuSC are also expressed by FAPs (97.1%, n=171) (Figure S3B, top panel). The list includes genes coding for the ECM components we analyzed with immunofluorescence staining (Collagen I, III, IV, VI, XV, and HCMN2). Only 5 core matrisome genes were considerably enriched in MuSCs compared to FAPs: *Lama3*, *Col19a1*, *Thsd4*, *Aebp1* and *Crim1* (Figure S3B, top panel and Table S2). In contrast, no core-matrisome gene (among the 139 genes analyzed) was enriched in gMyonuclei compared to MuSC. They were either not differentially expressed between the 2 populations (83.5%) or enriched in MuSCs (16.5%) (Figure S3B, middle panel). This suggested that MuSCs are a major source of matrisome genes or at least as important as the myofibers at the single-nucleus scale.

To identify genes enriched in MuSC compared to all 3 neighbors and define a MuSC matrisome enrichment signature we combined results of these 3 pairwise analyses to build matrisome enrichment diagrams for the core matrisome genes and the matrisome associated genes separately (Figure 4A). We found that *Col19a1*, *Aebp1* (Adipocyte enhancer-binding protein 1) and *Thsd4* (Thrombospondin Type 1 Domain Containing 4) were enriched in MuSCs compared to FAPs, endothelial cells and myofibers and thus representing the MuSC core matrisome signature. THSD4 protein is also known ADAMTS-like protein 6 and can regulate fibrillin matrix assembly (Tsutsui et al., 2010), whereas AEBP1 protein was shown to act as a regulator of fibrillar collagen assembly (Blackburn et al., 2018). In addition, the BM gene *Lama3* (laminin α3 chain) was enriched in MuSCs compared to both FAPs and myofibers, which is associated interstitial ECM.

**Figure 4.**
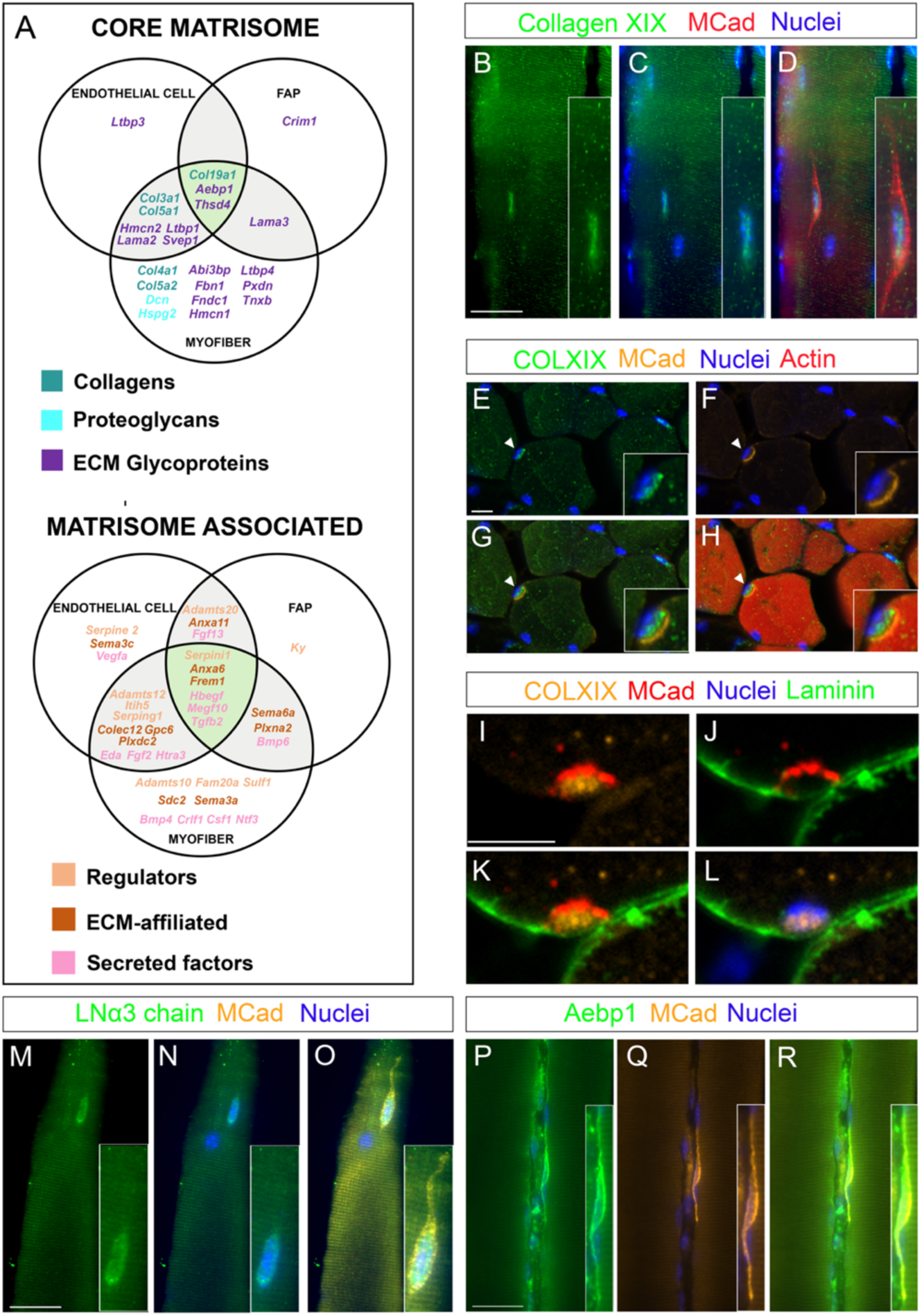
MuSCs display a matrisome signature distinct from the neighboring cell types. (A) Matrisome enrichment signature diagram showing matrisome genes whose expression is significantly enriched (log2 FC ζ 0.6) in MuSC compared to neighboring cells (endothelial cell, FAP, myofiber). Genes located in white areas are enriched in MuSC compared to one cell type only. Genes located in intersecting grey areas are enriched in MuSC compared to the two intersecting cell types. Genes located in the core green central area are enriched in MuSC compared to all three neighboring cell types and constitute the MuSC matrisome signature genes. (B-R) Co-immunostainings of mice muscle bundles (6-10 weeks old) with anti-M cadherin (Mcad) to label MuSCs (in orange or red as indicated) and Collagen XIX (B-L), LNα3 chain (M-O) or Aebp1 (P-R) antibodies (all in green). Nuclei were labelled with Hoechst (blue). All images are maximum Z-projections of a lightsheet Z-stack. Scale bars = 30 μm. (E-L) Co-immunostaining of muscle cryosections with (E,H) Collagen XIX (green) and MCad (orange) antibodies, (Actin is in red, pahalloidin staining) or (I,L) Collagen XIX (orange), MCad (red) and Laminin (green).White arrowhead points at the MuSC which are zoomed in the bottom right corner inserts (E,H). Nuclei were stained with Hoechst (blue). Scale bars = 10 μm. Images are maximum Z-projections of confocal Z stack. Muscles are from 8 weeks mice.

Matrisome-associated gene expression comparisons identified 6 genes that were specifically enriched in MuSCs and constitute the MuSC matrisome-associated signature: the regulator *Serpini1* (Neuroserpin), 2 ECM-affiliated proteins (*Frem1*, *Anxa6)* and 3 secreted factors (*Tgfb2, Megf10, Hbegf).* The couple *Sema6a-Plxna2* was also enriched in MuSCs compared to both FAPs and myofibers and could also specify MuSC niche. As we identified Collagen XIX, Aebp1 and Lama3 as potential new structural components of MuSC niche, we next characterized the presence of these ECM components in the MuSCs niche.

In the absence of specific antibodies against mouse collagen XIX, we generated antibodies targeting 2 peptides in the N-terminal and C-terminal parts of collagen XIX (Figure S4A). Immunostainings with Collagen XIX antibodies showed a strong signal at the level of the neuromuscular junction (NMJ) (Figure S4B-D and S4H-O) that was not observed in controls (Figure S4E-G). This is consistent with strong expression of *Col19a1* by a cluster of specialized myonuclei at the NMJ (as shown by UMAPs gene expression of *Col19a1* in 3 weeks and 5 months mouse Tibialis anterior snRNAseq available at the Myoatlas https://research.cchmc.org/myoatlas/). No Collagen XIX immunoreactivity was detected at the myofiber interfaces.

In contrast, Collagen XIX was observed as a strong intracellular signal in MuSCs (Figure 4B-D). Co-staining with pan-laminin antibodies that delineate the muscle BM confirmed the intracellular staining of Collagen XIX (Figure 4I-L). Likewise, LNα3 chain signal was not present at myofiber interfaces but showed a strong signal at the level of MuSCs cell body: 83% of MCad^+^ MuSCs were LNα3^+^ (Figure 4M-O) (n_MuSCs = 53; n_bundles = 7; n_muscles = 2; n_mice = 2) and 70% of GFP^+^ MuSC were LNα3^+^ (Figure S4E-G) (n_MuSCs = 61; n_bundles = 5; n_muscles = 2; n_mice = 2). Consistent with the enrichment of *lama3* expression in endothelial cells compared to MuSC (Table S2, MuSC/Endothelial cell Log2FC = −0.71), LNα3 chain signal also strongly lined up the contour of blood vessels (Figure S5A). In contrast, Aebp1 lined up the contour of MuSC cells, co-localizing with M-Cadherin signal (Figure 4P-R). Quantification showed that 83% of MCad^+^ MuSCs are Aebp1^+^ (n_cells = 71; n_bundles = 4; n_muscles = 2; n_mice = 2). The specific staining of Aebp1 delineating MuSC cells was also confirmed using a transgenic line expressing membrane-GFP under a Pax7-driven Cre-recombinase (Pax7-CreERT2; R26^mTmG^) to label MuSC (Figure S5B-D; 49% of GFP^+^ MuSCs displayed Aebp1 labeling (n_MuSCs = 65; n_bundles = 8; n_muscles = 3; n_mice = 3).

All together, these data revealed a MuSC matrisome gene signature including the specific expression of BMZ genes (*Col19a1, Lama3 and Frem1*) and regulators of ECM assembly (*Thsd4, Aebp1*) and identified COLXIX, AEBP1 and LAMA3 as new markers of MuSCs. This suggests that MuSC niche is chiefly produced and remodeled in a cell-autonomous manner.

### Matrisome of activated MuSCs displays features of cell motility and dysregulation of the qMuSC matrisome signature

In a previous study (Machado et al., 2017), RNA sequencing of qMuSCs was performed in parallel to the so-called early-activated MuSCs (aMuSCs), making possible to investigate the matrisome gene changes upon quiescence exit. aMuSCs display the classical transcriptional markers of early activated MuSCs including a strong induction of *Myod1* accompanied by significant drop of *Pax7* and Notch signaling targets (*Hes1* and *HeyL*) (Machado et al., 2017). We therefore used these transcriptomic data to perform a differential matrisome gene expression analysis between activated and quiescent states (Figure 5 and Table S3).

**Figure 5.**
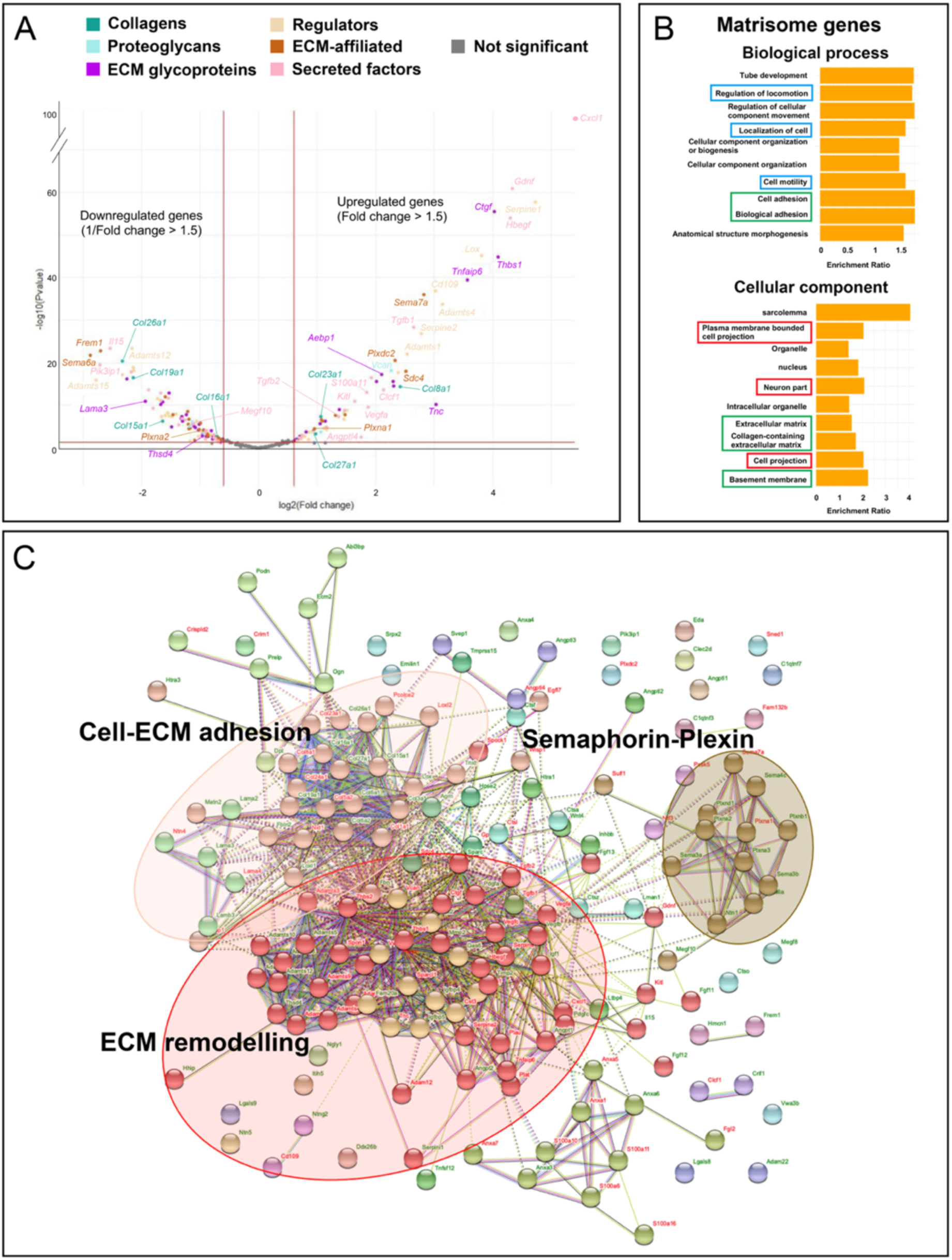
Changes in matrisome gene expression related to MuSC activation. (A) Volcano plot representing the differential expression of matrisome genes between qMuSC and aMuSC. Data points are colored according to each matrisome category. Red lines delimitate the zone where genes are considered as not differentially expressed (grey dots, |log2 FC| < 0.6 between vertical lines or p-values > 0.05 under the horizontal line). (B) Gene ontology enrichment analysis of Matrisome genes dysregulated upon MuSCs activation. Cellular component (bottom panel) or Biological process (top panel) domains were analyzed. Only Top 10 enriched GO terms are shown. GO terms linked to “Cell motility”, “Structural ECM” and “Neuron-related morphology” are highlighted in blue, green and red respectively. (C) Protein-protein interaction network analysis of the dysregulated matrisome genes upon MuSC activation produced with STRING database. The 169 dysregulated matrisome genes were input into STRING database and achieved a network of 169 nodes and 950 edges, with an enrichment p-values < 10-16. Clusters were obtained into STRING database with an MCL inflation parameter of 4. Genes are represented by colored bubbles. Each bubble color represents one cluster. Names of upregulated and downregulated genes are in red and green respectively. We highlighted 3 hubs of interests: Cell-ECM adhesion (salmon); ECM remodeling (red); Semaphorin-Plexin (brown).

First, this analysis revealed that genes identified as a signature of qMuSCs such as *Lama3, Col19a1*, *Thsd4*, *Frem1, Anxa 6, Serpini1 and Megf10* are all strongly downregulated in aMuSCs (Figure 5A and Table S3). In addition to *Col19a1 and Lama3*, another BMZ component *Col15a1* is also downregulated suggesting that aMuSCs remodel the BMZ to potentially favor motility. Consistent to this observation, loss of Collagen XV was shown to compromise the BMZ architecture and to favor epithelial tumoral cell invasion (Clementz et al., 2013). Strikingly, the chemokine *Cxcl1*, which is a potent inducer of neutrophil chemotaxis (Yang et al., 2019; Kroeze et al., 2012), was instead the most upregulated gene with a 151-fold change. Along this line, the couple *Sema6a/Plxna2* was strongly downregulated while the couple *Sema7a/Plxna1* was upregulated upon MuSC activation. Anti-migratory effects on lung cancer cells were attributed to semaphorin 6A (Chen et al., 2019) while Semaphorin 7A promoted immune cells migration (Morote-Garcia et al., 2012; van Rijn et al., 2016) suggesting that this switch in semaphorin activity could also favor motility.

Accordingly, GO analysis showed a strong enrichment in gene sets involved on one hand in “extracellular matrix”, basement membrane” and “collagen-containing extracellular matrix and on the other hand in the “regulation of locomotion” “localization of cell” and “cell motility”” (Figure 5B). In accordance with these data, STRING analysis of the dysregulated matrisome genes revealed 3 major hubs that were named according to the gene products involved in the protein-protein networks (Figure 5C): (1) the “ECM remodeling” hub made of about 50 genes among which numerous genes encoding matrix metalloproteases (members of the ADAMTS and MMP families); (2) the cell-matrix adhesion hub and (3) the so-called semaphorin-plexin hub showing a striking downregulation of all Semaphorine-Plexin genes. The expression of “neurogenesis related” genes was an important feature of qMuSC matrisome. GO analysis showed enrichments in gene sets associated with neuron biology (i.e the cellular component terms ““Neural part” and “cell projections”) (Figure 5B).

All together these data revealed that aMuSCs downregulate the matrisome signature and express a matrisome gene set that should favor the breakdown of BMZ and the remodeling of the surrounding ECM to promote cellular migratory behavior. These data suggest that qMuSCs deliberately leave the niche, aligning with their necessity to relocate and repair injured muscle in vivo.

### Comparative analysis of muscle dissociation and muscle injury redefines the MuSC early activation ECM profile

We found that activated MuSCs by tissue dissociation downregulated the quiescent MuSC matrisome genes and induced a matrisome favoring cell motility. To test whether these findings hold true in a more physiological context we used snRNAseq data from injured muscles (Machado et al., 2021). We previously reported that MuSC activated *in vivo* by muscle injury or *ex vivo* by tissue dissociation show similar transcriptional kinetics, at least for the five major gene clusters described (Machado et al., 2021). Here, we extracted matrisomes from intact and injured MuSC nuclei and performed differential expression analysis between these 2 conditions (Table S3). To be consistent with our previous analysis (Figure 5), we considered genes to be expressed by MuSC only when the average expression was above 0.026 for at least one condition, and genes to be dysregulated only when |log2FC| ζ 0.6 with a p-value < 0.07 between 2 conditions. We then compared the dysregulation status of matrisome genes upon tissue dissociation versus tissue injury (Figure 6A, Table S3). First, almost 75% of the matrisome genes that were dysregulated upon tissue dissociation were not significantly dysregulated upon tissue injury (only 41 matrisome genes were dysregulated compared to 163 upon tissue dissociation). Interestingly, the matrisome features we pointed out as favoring cell motility, such as the massive upregulation of *Cxcl1* or downregulation of *Sema7a*, were not observed upon tissue injury. Similarly, expression of semaphorin/plexin genes expression was not downregulated upon MuSC activation *in vivo* with the exception of the couple *Sema6a/Plxna2*. We also noticed that 14 ECM regulators, including several members of the ADAMTS family, *Cd109* and *Serpini1* are among the 41 matrisome genes dysregulated in MuSC upon muscle injury and that ECM regulators represent no less than a third of matrisome genes similarly dysregulated upon tissue dissociation and tissue injury (9/28 genes). This suggests that MuSCs activated *in vivo* after muscle injury also extensively remodel the surrounding ECM. Along this line, the BMZ components *Lama2, Col8a1, Col5a2 and Col19a1* were similarly dysregulated in tissue dissociation and tissue injury.

**Figure 6.**
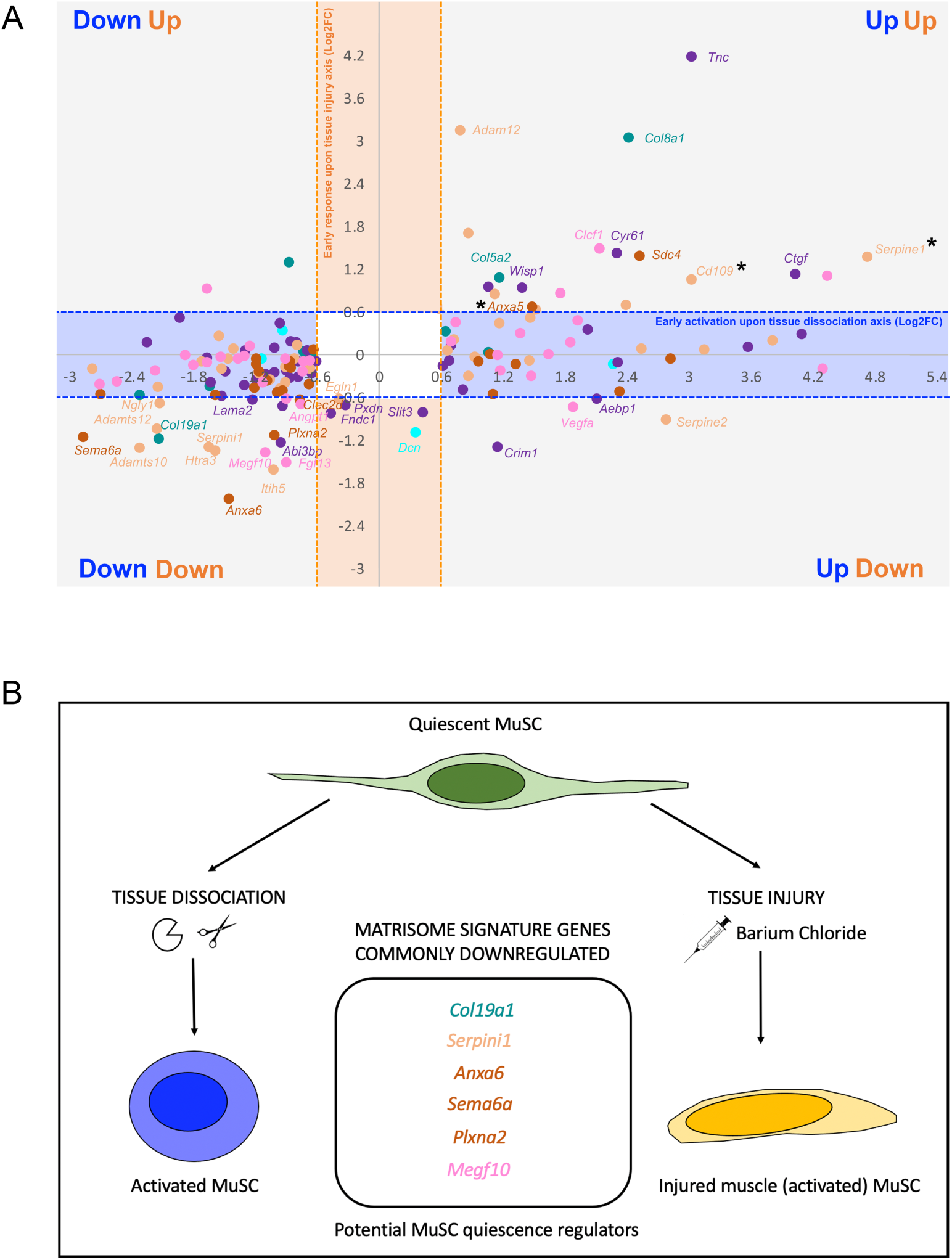
Differences and similarities between matrisome gene expression in MuSC after tissue dissociation and tissue injury. (A) Comparison of Matrisome dysregulated genes between “Early activation upon tissue dissociation (aMuSC/qMuSC) and “Early response upon muscle injury” (Injured muscle MuSC /Intact muscle MuSC). Each dot corresponds to a gene. x coordinates correspond to the log2FC of the expression level between quiescent and activated MuSC upon tissue dissociation (blue axis). Y coordinates correspond to the log2FC of the expression level between MuSC from an injured versus an intact muscle (orange axis). Only genes significantly dysregulated in at least one situation are present in the graph. Labeled genes in the grey zones are significantly dysregulated in both situations. Labeled genes in the orange zone are significantly dysregulated upon muscle injury but not upon tissue dissociation. Unlabeled genes are significantly dysregulated upon tissue dissociation but not muscle injury. Unlabeled genes (in grey zones and blue zones) are significantly dysregulated upon tissue dissociation but not muscle injury. Genes with stars have a p value for log2FC (Injured muscle MuSC/Intact muscle MuSC) between 0.05 and 0.07. (B) Schematic summarizing the MuSC matrisome signature genes similarly regulated upon tissue dissociation and tissue injury.

Regarding the qMuSC matrisome signature, we found that *Col19a1, Serpini1, Anxa6, Sema6, Plxna2* and Megf10 are downregulated both upon tissue dissociation and tissue injury. Combined with the fact that these 6 genes are all part of the top 50 matrisome genes expressed by qMuSC, this suggests that they might be new regulators of MuSC quiescence (Figure 6B). Notwithstanding differences in the depth of sequencing and in the length of activation, altogether our results indicate that the response of MuSC matrisome-related genes differs depending on the activation context, however the ECM downregulation of the qMuSC-specific matrisome genes is conserved during *ex-vivo* and *in vivo* MuSC activation.

## DISCUSSION

There is growing evidence showing that quiescent cells are transcriptionally and metabolically active (van Velthoven and Rando, 2019; Roche et al., 2017). Based on the transcriptomic data used for this study, murine qMuSC expressed about 17% of their genome, which is similar to what aMuSCs expressed upon activation (about 15%). Interestingly, similar counts were also seen for the transcriptional activity of qMuSC and aMuSC at the matrisome level (about 22% for both) indicating the key role of ECM components in MuSC behavior. In support to these data, the MuSC matrisome was found to be as rich and diverse as that of the FAPs (Theret et al., 2021).

Our bioinformatics analysis revealed that qMuSC are nested in an environment particularly rich in collagens and ECM glycoproteins whose primary function is to modulate tissue mechanical properties and cell behavior, respectively. We showed that MuSCs are in close proximity not only with BMZ collagens (i.e. collagen IV, VI, XV) but also interstitial collagens (i.e. collagen I, III), raising the question whether MuSCs actively contribute to the assembly this collagen-rich matrix around them. Supporting this idea, we show that qMuSC strongly express *Aebp1* that specifically lined up their contour. Aebp1 is a carboxypeptidase that binds to collagens through its Discoidin domain and positively regulates collagen I fibrillogenesis (Blackburn et al., 2018), suggesting that MuSCs could locally assemble collagen I fibrils through AEBP1 activity. Interestingly, *Aebp1* is downregulated in MuSCs after muscle injury but not during tissue dissociation with collagenase treatment. This suggests that MuSCs might remodel their collagen I matrix to seclude themselves from the niche upon *in vivo* activation. Consistent with this, *Ddr2,* a tyrosine kinase collagen receptor that was shown to inhibit collagen I fibrillogenesis (Mihai et al., 2006; Flynn et al., 2010) is strongly upregulated in MuSC upon activation both in muscle dissociation and muscle injury (logFC intact/injured muscle MuSC=1.99; logFC aMuSC/qMuSC=2.8; non-matrisome genes analysis, data not shown).

Bioinformatics analysis revealed that a significant proportion of the matrisome genes expressed by qMuSC are basement membrane tool-kit genes (namely collagen IV, laminins, perlecan and nidogens) and genes involved in the organization of the BM, such as *Frem*. qMuSC actively express BM tool-kit genes and show specificity in expressing at the transcriptional and protein levels *Lama3* and the basement membrane associated gene *Col19a1,* indicating that Collagen XIX and LNα3 chain are new MuSC markers. This local BM specificity may act as a niche that favors MuSC quiescence. *Col19a1* and *Lama3* were both downregulated in aMuSC upon tissue dissociation suggesting their role in maintaining MuSC quiescence. Additionally, the aMuSC transcriptome is enriched in genes involved in ECM degradation, cell migration, and motility, consistent with early activation. Notably, Cxcl1, involved in cell motility (Yang et al, 2019), and Anxa1, which promotes MuSC migration and differentiation post-injury (Bizzaro et al. 2012), are upregulated in aMuSCs. Many genes dysregulated in activated MuSCs during tissue dissociation were not significantly dysregulated in MuSCs following muscle injury indicating that, *in vivo,* activation is influenced by specific interactions and mechanical contexts that finely tune the MuSC response.

Intriguingly, qMuSCs express genes involved in neurogenesis, which are significantly dysregulated upon MuSC activation. Our GO analysis suggests that these genes could be involved in establishing and remodeling long cell projections, typical of neuron morphology. Our quantitative morphological analysis showed that 81% of MuSCs display neuron-like morphology with long protrusions. Expression of matrisome genes linked to cell protrusions was significantly dysregulated upon activation, consistent with the observation that MuSCs lose their neuron-like projections upon activation.

*Serpini1* and *Abi3bp*, top expressed genes in qMuSCs, are significantly downregulated upon activation in both tissue dissociation and tissue injury. These genes may play a role in shaping MuSC morphology via cytoskeletal regulation, stabilizing MuSC quiescence. *Serpini1* encodes for Neuroserpin which was involved in shaping dendritic protrusions in cultured neurons (Borges et al., 2010) and *Abi3bp* encodes for the ECM glycoprotein “ABI Family Member 3 Binding Protein” which has been shown to restrict dendritic branching and outgrowth of interneurons (Cheng et al., 2009).

Quiescent MuSCs also express Agrin, a protein essential for neuromuscular junction formation (Daniels, 2012). MuSCs were shown to contribute to neuromuscular junction integrity and regeneration post-denervation (Liu et al., 2015; Liu et al., 2017; Larouche et al., 2021). Furthermore, the qMuSC matrisome signature gene *Col19a1* identified in our study, contributes to synaptic contact formation in hippocampal neurons and shapes neuromuscular junctions (Su et al., 2010). Given the enrichment of qMuSCs matrisome genes related to neuron morphology and differentiation, it is plausible that MuSCs play a role in motor axonogenesis and axon guidance. When activated, MuSCs upregulate genes involved in neuron homeostasis and repair, such as Wisp1, Cyr61, Clcf1, and TnC. WISP1 promotes neuronal repair (Maiese, 2015) and Cyr61 Schwann cell proliferation (Cheng et al., 2021). CLCF1 sustains embryonic motoneurons in vitro and promotes astrocyte differentiation, and its expression increases after nerve injury (Crisponi et al., 2022). Upregulation of Tenascin C, a motoneuron guidance cue, is common in neural injury models. Recent single-cell analysis beautifully highlighted the cross-talk between MuSCs and other resident cells during muscle regeneration (De Micheli et al.,2020).

In conclusion, this study revealed specific matrisome gene signatures of quiescent and activated MuSCs that give insights in gene sets associated to their respective behavior and the possible contribution of muscle residing cells in building the niche by producing ECM components and remodeling the surrounding ECM. As such, the data provided interesting gene candidates to target for muscle repair and the treatment of degenerative muscle disease as well as to design artificial niche for tissue engineering.

## METHODS

### RNAseq dataset availability

The NGS RNAseq and single nuclei datasets used in this study had been published previously (Machado et al., 2017; Machado et al., 2021) and are available online at the Gene Expression Omnibus website (https://www.ncbi.nlm.nih.gov/geo/) under the GEO accession numbers GES103162 and GSE163856 respectively.

### NGS RNAseq data analysis

In order to highlight a subset of genes that could play key biological functions in MuSC quiescence and activation, we decided to reduce the size of the analyzed dataset by setting an expression cut-off at 1000 reads at the quiescent state (q MuSC, refered as T0-SC in Machado et al., 2017): only the genes above this expression level were considered as expressed in our study. For the differential expression analysis between activated MuSC (aMuSC, refered as T3-SC in Machado et al., 2017) and qMuSC, genes under this 1000 reads cut-off in both quiescent and activated states were discarded from the analysis. Identification and classification of matrisome genes present in the dataset were performed using the online Matrisome Annotator tool (MIT, Matrisome project, http://matrisomeproject.mit.edu/) (Naba et al., 2012). Count data from quiescent and activated MuSCs were analyzed with R software version 4.1.0 with the Bioconductor DESeq2 package version 1.32.0 (Love et al., 2014). Normalization and dispersion estimation were performed with DESeq2 using the default parameters. A statistical test for differential expression was performed using a generalized linear model to test for differential expression between qMuSC and aMuSC. For each pairwise comparison, raw p-values were adjusted for multiple testing according to the Benjamini and Hochberg (BH) procedure (Benjamini and Hochberg, 1995) and genes with adjusted p-values < 0.05 and a | log2foldchange | > 0.6 were considered as differentially expressed. The resulting volcano plot in Figure 5A was generated using the ggplot2 R package.

### Single nuclei RNAseq data analysis

The Seurat object containing single nuclei clustered data from intact muscles, injured muscles, and digested muscles (kindly provided by Leo Machado, Relaix laboratory, UPEC faculty of medicine, France), have been reanalyzed using R software version 4.1.0 with the package Seurat version 4.0.3. To be consistent with the reads cut-off value applied to the qMuSC analysis, a gene expression cut-off value of 0.026 was applied to these data sets.

### Matrisome signature analysis

For the matrisome signature analysis, we first extracted the intact muscle data with the “subset” and “createSeuratobject’’ functions. This new Seurat object created with only intact muscle nuclei data has been reimported on R software and a differential expression analysis has been conducted between MuSCs nuclei and FAP nuclei, MuSCs nuclei and Myonuclei, or MuSC nuclei and Endothelial cell nuclei on the 22161 genes retrieved in this dataset. This differential expression between the clusters has been conducted using the “findmarkers” function, and a student t-test has been used to find significant differences between the cell types. Genes with a p value < 0.05 and a | log2Foldchange | > 0.6 were considered differentially expressed between the nuclei clusters. A gene expression cut-off to define genes as “expressed genes” in our study was empirically determined at 0.026 in order to relatively match the 1000 reads cut-off used in the NGS RNA seq analysis. When performing the FAP/MuSC, Myofiber/MuSC and Endothelial cell/MuSC pairwise analyses, genes under the 0.026 threshold for both clusters of the pair analyzed were discarded. Identification and classification of matrisome genes present in this dataset were performed as described above for NGS RNAseq data. The resulting volcano plots in Figure S3 were generated using the ggplot2 and ggrepel R packages.

### Injured vs intact muscle MuSC matrisome analysis

For this comparative analysis, we first extracted the intact muscle and injured muscle data with the “subset” and “createSeuratobject’’ functions. This new Seurat object has been reimported on R software and a differential expression analysis has been conducted between MuSCs nuclei from intact muscle versus injured muscle using the “findmarkers” function, and a student t-test has been used to find significant differences between the two conditions. Genes with a p value < 0.07 and a | log2Foldchange | > 0.6 were considered differentially expressed between the nuclei clusters. We used the same expression cut-off than for the matrisome signature analysis and genes under the 0.026 threshold in both conditions were discarded. Identification and classification of matrisome genes present in this dataset were performed as described above for NGS RNAseq data. In order to highlight matrisome markers of qMuSC and/or a subset of genes that could play key biological functions in MuSC quiescence, the dataset was pre-filtered by setting a gene expression cut-off at 1000 reads so that only the genes above this expression level are considered in the following analysis.

### Gene ontology analysis

The gene set ontology enrichment analysis has been performed using the default parameters in WebGestalt (WEB-based Gene SeT AnaLysis Toolkit) online tool (http://www.webgestalt.org/), either comparing the list of the matrisome expressed genes against all the matrisome genes of the mouse genome (analyses in Figure 2A, 2C, 5B), or comparing all expressed genes minus the matrisome expressed genes against all mouse genome minus the matrisome genes (analyses in Figure 2D). Gene sets with an FDR < 0.05 were considered enriched, where the FDR is the estimated probability that a gene set with a given normalized enrichment score represents a false positive finding. The results were plotted using the R package ggplot2.

### Functional interaction network analysis (STRING)

In order to perform a functional interaction analysis of the dysregulated matrisome upon MuSC activation (Figure 5C) and to further characterize the basement membrane gene network expressed by quiescent MuSC (Figure 2B), we used the online STRING database (https://string-db.org/) to build functional networks. This database provides information on protein functional interactions based on information from high-throughput experimental data as well as database and literature mining, and predictions based on genomic context analysis. The STRING database gives an association score for two significantly functionally interacting proteins. This score is calculated using various parameters such as the neighborhood score, fusion, co-occurrence, homology, co-expression, experimental score, database score and text mining score, among others. A higher score indicates greater strength of functional association and interaction between proteins. The chains also provide network analysis tools such as clustering. In the current study, the clustering based on Markov Cluster Algorithm (or MCL clustering) provided in STRING version 11.5 was used to cluster the genes. The choice in the MCL inflation parameter value to perform the clustering (MCL= 2 for Figure 5C and MCL=4 for Figure 2B) was empirically determined to tune the appropriate level of stringency to reveal biologically relevant clusters. The downloaded STRING result in Figure 5C has been manually edited to color code genes name (upregulated vs downregulated) using Adobe illustrator software.

### Mice muscle samples

Wild-type tibialis anterior muscles for this study were obtained from 6 to 12 weeks old C57BL/6 wild-type male or female mice. Those mice were from Jackson laboratories and were handled at PEHR facility (Institut de Génomique Fonctionnelle de Lyon, Lyon, France) according to French and European Union guidelines. Fixed tibialis anterior from Tg : TgPax7-CreRT2 ;R26^mTmG^ 8 weeks mice (2H fixation at RT in 2% PFA in 1X PBS and storage until use in 1X PBS) were obtained from Philippos Mourikis research group (Relaix Laboratory, IMRB, France).

### Production and purification of antibodies against mouse Collagen XIX

Polyclonal antibodies specifically recognizing mouse Collagen XIX were raised in guinea pigs by co-immunization with the following synthetic peptides: CPTLRTERYQDDRNKS and PEDCLYPAPPHQQAGGK located in the non-collagenous domain NC6 and NC1 respectively (see Fig. S3 for details). Specific antibodies to Collagen XIX were then purified by antigen affinity chromatography against a mix of both peptides. Peptide synthesis antibody production, and purification were provided by Covalab (Bron, France).

### Immunofluorescence staining

For whole-mount immunostainings, tibialis anterior (TA) muscles were sampled from mice, fixed 2H at RT in 2% PFA in 1X PBS and quickly rinsed in 1X PBS before overnight storage at 4 degrees in 1X PBS. Muscles were dissected to isolate fibers bundles whose size was ranging from 3 to 15 fibers. Bundles were then permeabilized for 1H at RT in 0.5% Triton X-100 in 1X PBS and incubated 3-4H at RT in blocking solution (20% Donkey serum,0.1%Triton X-100 in 1X PBS). Bundles were incubated at 4°C for 22 - 23 H with primary antibodies diluted in 5% donkey serum, 0.1% Triton X-100 in 1X PBS. After being washed 4 x 20 min in 0.1% Triton X-100 in 1X PBS, bundles were incubated at 4°C for 22-23 H with secondary antibodies and phalloidin or alpha-bungarotoxin diluted in 5% donkey serum, 0.1% Triton X-100 in 1X PBS. Upon removal of secondary antibodies, nuclei were stained with Hoechst 33342 (SIGMA 2.5 μg /ml final in 0.1% Triton X-100 in 1X PBS) for 15 min at RT, and bundles were washed 3 x 20 min in 0.1% Triton in 1X PBS before being transferred in 1X PBS for subsequent mounting. All washes were performed at RT.

For immunofluorescence on frozen tissue sections, TA muscles were fixed overnight at 4°C in 4% PFA in 1X PBS, quickly rinsed in 1X PBS, embedded in OCT freezing medium, and then snap-frozen in cold isopentane. 20 μm cryosections were cut using a Leica cryostat and stored at −20°C until use. Sections were left 15 min at RT, rehydrated for 5 min in 1X PBS and incubated 1 hour at RT in blocking buffer (20 % Donkey serum in 0.1% Triton in 1X PBS). Primary antibodies were then incubated overnight at 4°C in a homemade wet chamber. After quick washes in 1X PBS (4x5 min), secondary antibodies or phalloidin drug were incubated for 1H at RT. After 2x 5 min washes in 1X PBS, nuclei were stained for 7 min at RTwith Hoechst 33342 (SIGMA 2.5 μg /ml final in 0.1% Triton X-100 in 1X PBS). Samples were washed 5 min in 1X PBS and mounted in Dako Fluorescent Mouting medium (Dako, S3023) and stored at 4°C until imaging. All washes were performed at RT.

Primary antibodies used in this study were : rabbit anti-human collagen I (Novotec 20111; 1/40); rabbit anti-rat collagen III (Novotec 20341; 1/100); rabbit anti-human collagen VI (Novotec 20611; 1/200); rabbit anti-human collagen IV (Novotec 20411; 1/20); rabbit anti-mouse collagen XV NC8 antibody kindly gifted by Takako Sazaki (Oita University, Japan; 1/1000); rabbit anti-human Hemicentin2 (novus biologicals NBP2.30512; 1/100); sheep anti-human M-cadherin (R&D Systems AFH096; 1/100); rat anti-mouse PDGFRA (Invitrogen 17-1401-81; 1/500); Purified polyclonal guinea pig anti-mouse Collagen XIX (see production of Collagen XIX antibodies in the methods section ; 1/100) ; rabbit anti-human Aebp1 (Origene TA329346 ; 1/50) ; rabbit unpurified antisera anti-mouse laminin alpha3AIIIa (1/500) ; rabbit anti-mouse Laminin (SIGMA L9393 ; 1/400) ; Chicken anti-GFP (Abcam ab13970 ; 1/1000). Secondary antibodies and drugs used in this study were : donkey anti-rabbit alexa fluor 488 (Invitrogen A21206 ); donkey anti-rabbit alexa fluor 647 (Invitrogen A31573) ; donkey anti-sheep alexa fluor 546 (Invitrogen A21098) ; donkey anti-sheep alexa fluor 647 (Invitrogen A21448) ; donkey anti-rat alexa fluor 488 (Invitrogen A21208) ; donkey anti-guinea pig alexa fluor 488 (Jackson immuno Research ref 706-545-148) ; donkey anti-guinea pig TRITC (Jackson immuno Research ref 706-025-148) ; goat anti-chicken alexa fluor 488 (Invitrogen A11039) alpha-bungarotoxin-TRITC (SIGMA T0195 resuspended at 1mg/ml in sterile water ; working dilution 1/100) ; Phalloidin-Alexa 647 (Invitrogen A22287 resuspended in 1.5ml MeOH according to manufacturer instructions ; working dilution 1/80). All secondary antibodies were used at a 1/500 dilution except for goat anti-chicken alexa fluor 488 used at 1/1000 dilution.

### Imaging and Image processing

Muscle bundles after whole-mount immunostaining were mounted in glass capillaries (1.4 mm diameter) in 1% agarose in 1X PBS and imaged with a Lightsheet Z.1 (ZEISS, 20x objective) available at the CIQLE facility (University Lyon1, Lyon, France). Immunostainings on cryosections and some muscle bundles after whole-mount immunostaining (for the MuSC morphology analysis, mounted in 50% glycerol in 1X PBS) were imaged at with an inverted confocal microscope Zeiss LSM 780 and a 20X objective (IGFL, Lyon, France). Image processing including 3D image rendering, muscle interfaces reconstructions and generation of virtual transverse sections was performed using Imaris software version 9.1 available at the PLATIM facility (ENS LYON, Lyon, France). ImageJ software was used to do maximum Z projections.

## ACKNOWLEDGEMENTS

We thank Damien Sery for his assistance with the dissection of mouse skeletal muscle and Bruno Chapuis (CIQLE, University of Lyon) for his valuable technical support with the light sheet microscope. This study was funded by an ANR grant (ANR-19-CE13-0010).

## ABBREVIATION LIST

aMuSC: Activated Muscle Stem Cell
BM: Basement membrane
BMZ: Basement membrane zone
ECM: Extracellular matrix
FAP: Fibro-adipogeno-progenitor
GO: Gene Ontology
MuSC: Muscle Stem Cell
qMuSC: Quiescent Muscle Stem Cell

## Supplementary Figures

**Figure S1 related to Figure 1.**
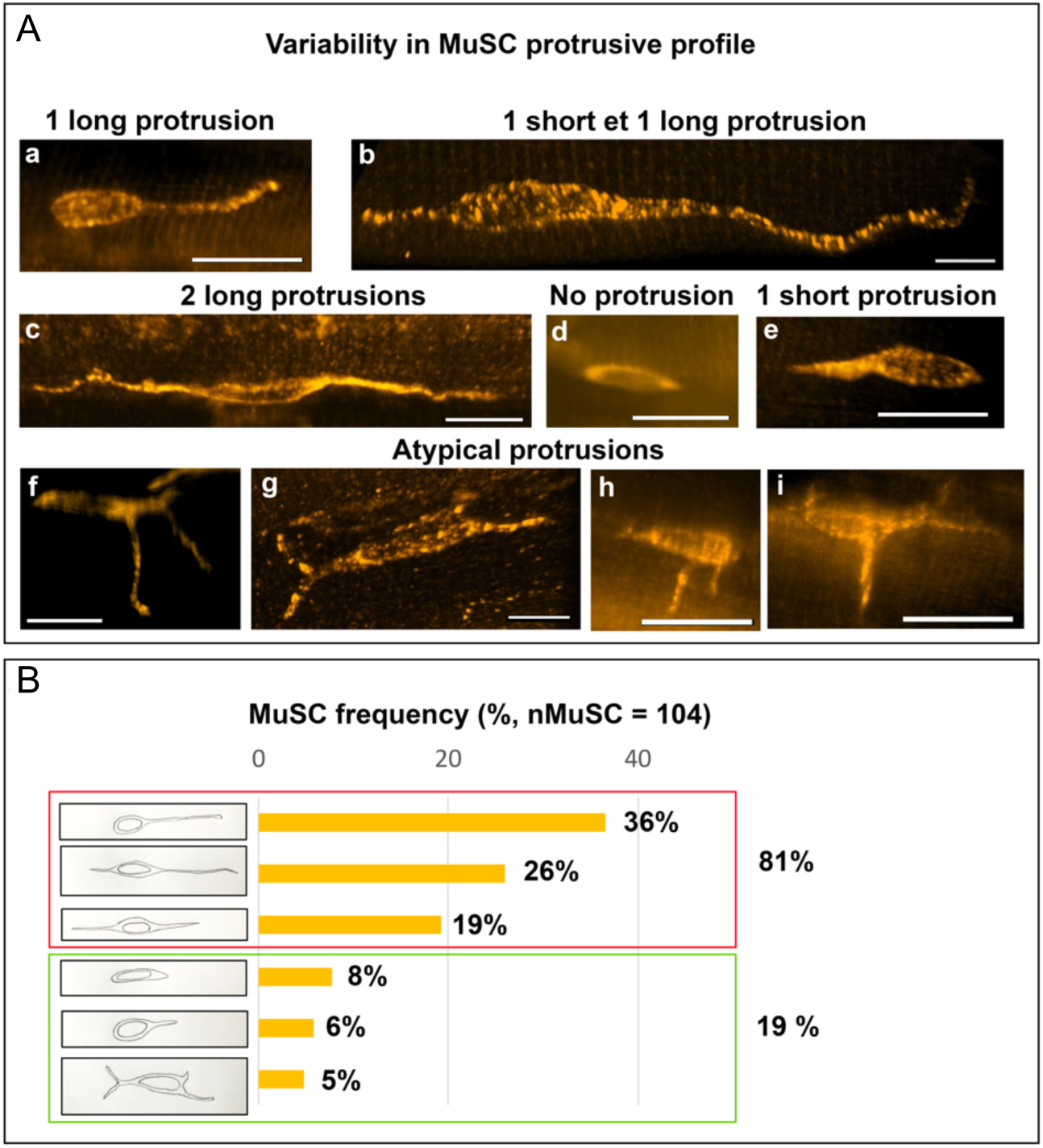
Analysis of MuSC morphology. (A) MuSCs exhibit different protrusive profiles in a muscle bundle. 3D projections of confocal (b,c,g) or lightsheet (a,d,e,f,h,i) imaging of MuSCs labeled with Mcad (orange) in a whole muscle bundle. Six protrusive profiles were defined based on the number, the length and the orientation of the protrusions. Long protrusion = protrusion whose length ≥ one cell body length. Short protrusion = protrusion whose length < one cell body length. Scale bars = 20 μm in a,d,e,h,i; 10 μm in c,g; 15 μm in f; 8 μm in b. (B) Quantitative analysis of MuSC protrusive profiles. Each drawing represents a protrusive profile as defined in (A). Pooled data from 13 muscle bundles (n muscles = 3; n mice = 2).

**Figure S2 related to Figure 3.**
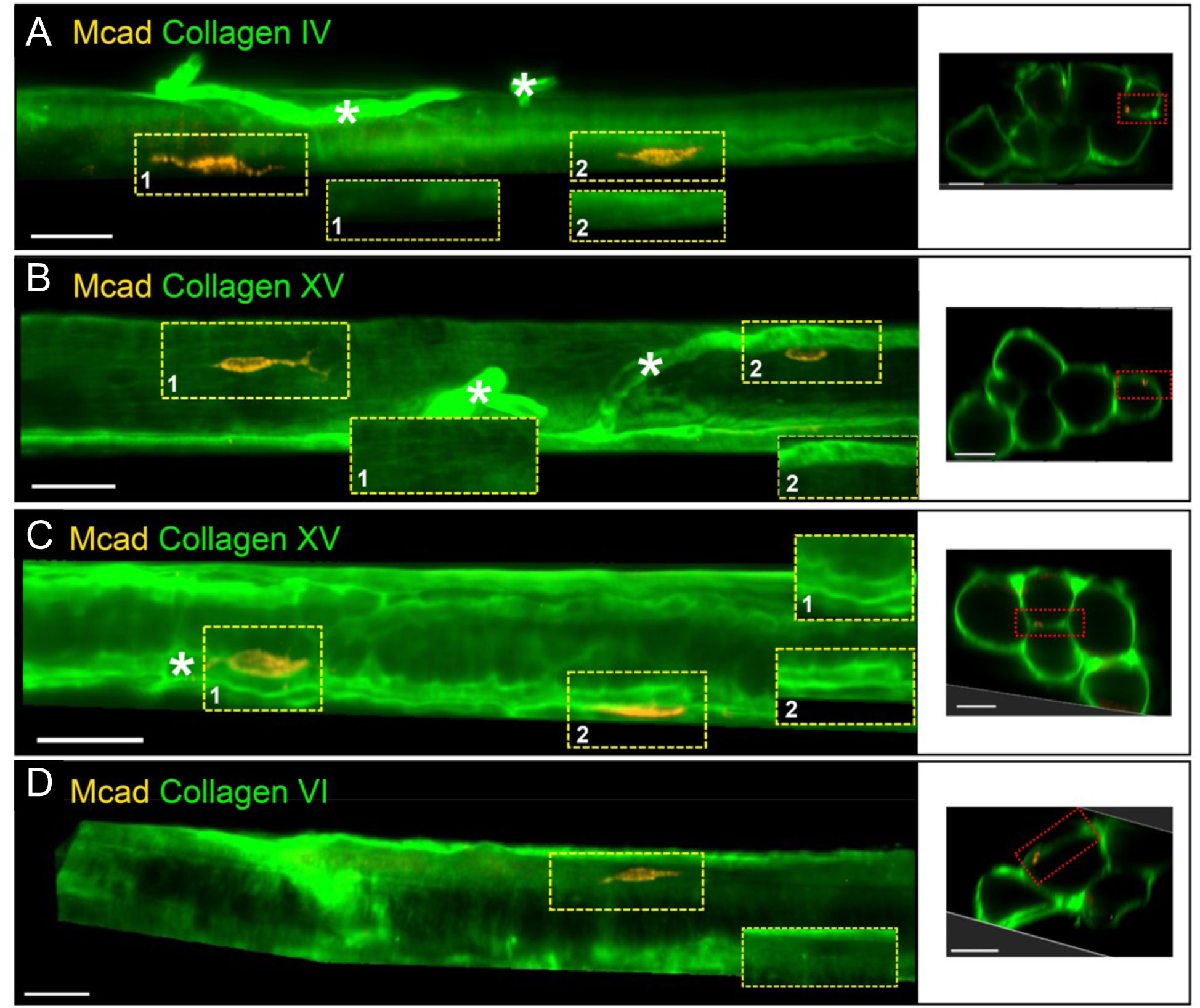
The BM/BMZ collagens I, XV or VI do not organize into a specific pattern around MuSC. (A-E) Co-immunostaining of mice muscle bundles (8-12 weeks old) with anti-M cadherin (Mcad) to label MuSCs in orange and Collagen IV (A), Collagen XV (B,C) or Collagen VI (D) antibodies (green). Projected top views of 3D myofiber interfaces. The corresponding transverse sections indicate the position of each interface in the whole muscle bundle (red rectangle). Cropped areas show regions in the corresponding yellow frames without MuSCs to visualize matrix signal at the cell position. Scale bars= 30 μm. White stars indicate blood vessels.

**Figure S3 related to Figure 4.**
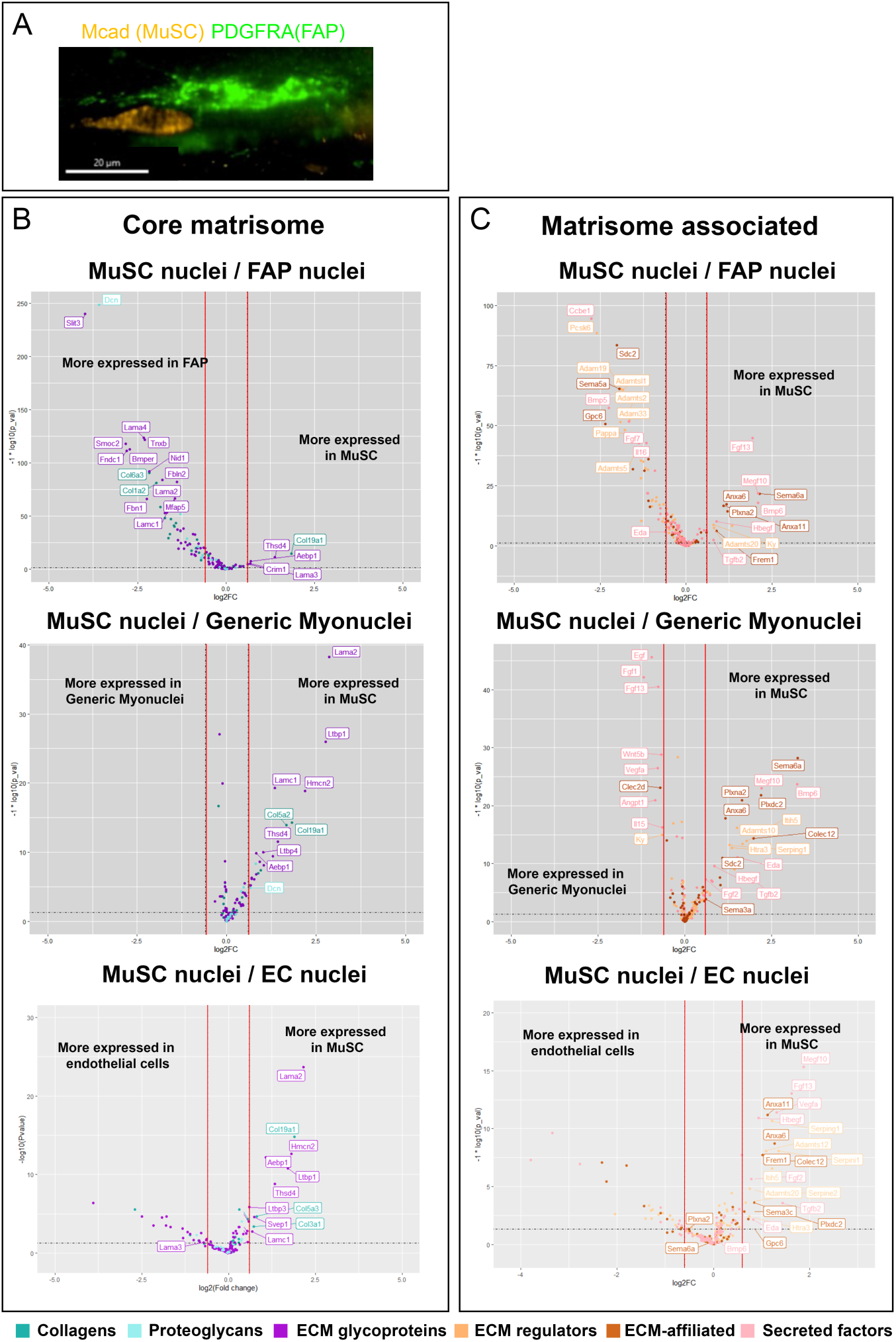
(A) MuSCs are in close proximity with FAPs. 3D projection of lightsheet imaging of MuSCs labeled with Mcad (orange) and FAPs labeled with PDGFRA (green) in a whole muscle bundle. Scale bar = 20 μm. (B and C) Volcano plots representing the difference in expression between FAP nuclei and MuSC nuclei (top panels), Myonuclei and MuSC nuclei (middle panels) or Endothelial cell (EC) nuclei and MuSC nuclei (bottom panels) for Core matrisome genes (B) or Matrisome associated genes (C). Data points are colored according to each matrisome category. Red lines delimitate the zone where genes are considered as not differentially expressed (|log2 FC| < 0.6 or p value > 0.5 under the grey dotted line).

**Figure S4 related to Figure 4.**
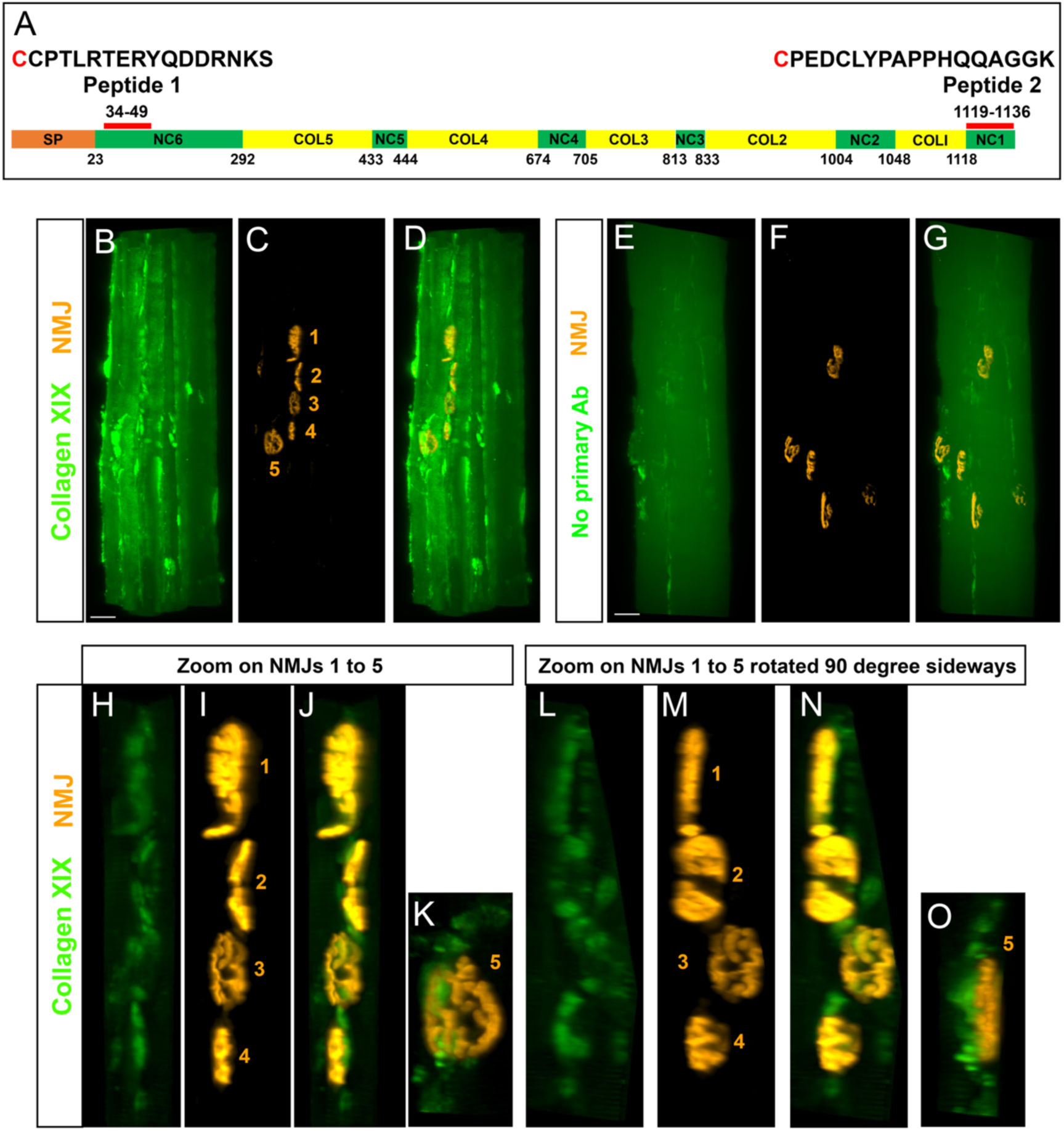
(A) Production of polyclonal antibodies against mouse collagen XIX. Schematic representing the primary sequence of the mouse Collagen XIX alpha chain (1136 aa) with collagenous (COL) and non-collagenous (NC) domains. SP=signal peptide. The first amino acid of each domain is indicated as well as the sequence and position of the two synthetic peptides which were co-injected into guinea pigs to generate antibodies. The initial cysteine (red) in the synthetic peptides is not part of collagen XIX sequence. (B-O) Collagen XIX is deposited at the neuromuscular junction (NMJ). 3D projections of lightsheet imaging of whole muscle bundles (6 weeks mice) co-labelled with alpha-bungarotoxin to visualize NMJs (orange) and Collagen XIX antibodies (green, B-D and H-O). Negative control without primary antibody is shown in (E-G). (H-O) are zoomed and cropped areas of B-D around the NMJs numbered in C with the same angle of view (H-K) or rotated 90 degrees sideways (L-O). Scale bars = 30 μm.

**Figure S5 related to Figure 4.**
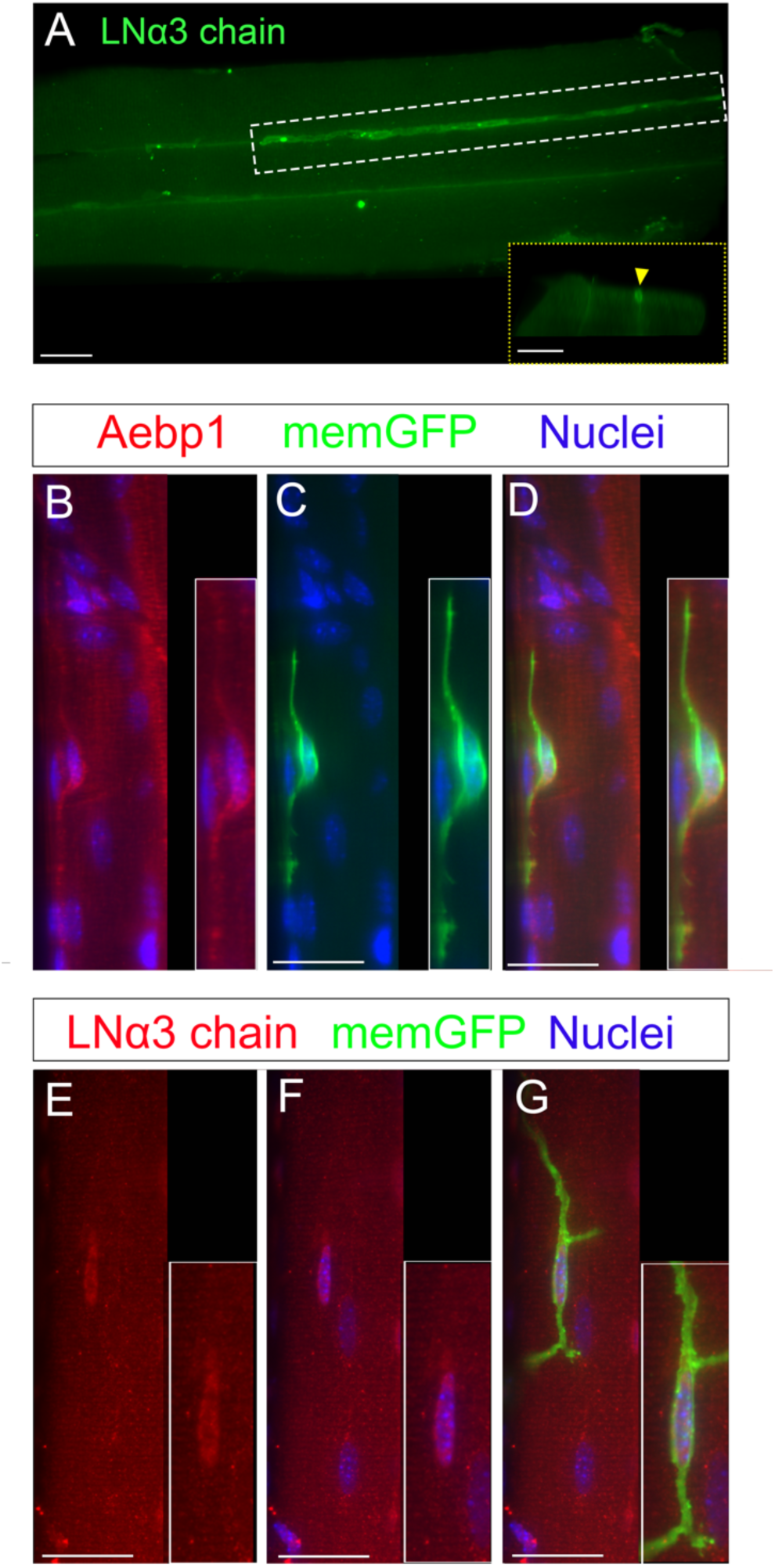
(A) Immunostaining of LNα3 chain on a wild-type muscle bundle (8 weeks old) showing LNα3 chain labelling on blood vessels (white arrow and white dotted box). Yellow dotted insert corresponds to virtual transverse section at the level of the blood vessel in the white dotted box (yellow arrowhead). (B-G) Co-immunostainings of transgenic mice (TgPax7-CreRT2;R26^mTmG^) muscle bundles (8 weeks old) with anti-GFP antibodies to label MuSCs in green (Pax7 expressing cells switch on the expression of a membrane-GFP (memGFP) upon tamoxifen treatment), and with Aebp1 (B-D) or LNα3 chain (D-F) antibodies (red). Nuclei were labelled with Hoechst (blue). Images are maximum Z-projections of a lightsheet Z-stack except in A which is a top view of 3D projection of the whole muscle bundle (also acquired with a lightsheet). Scale bars = 30 μm.

## Supplementary Tables

**Table S1.**
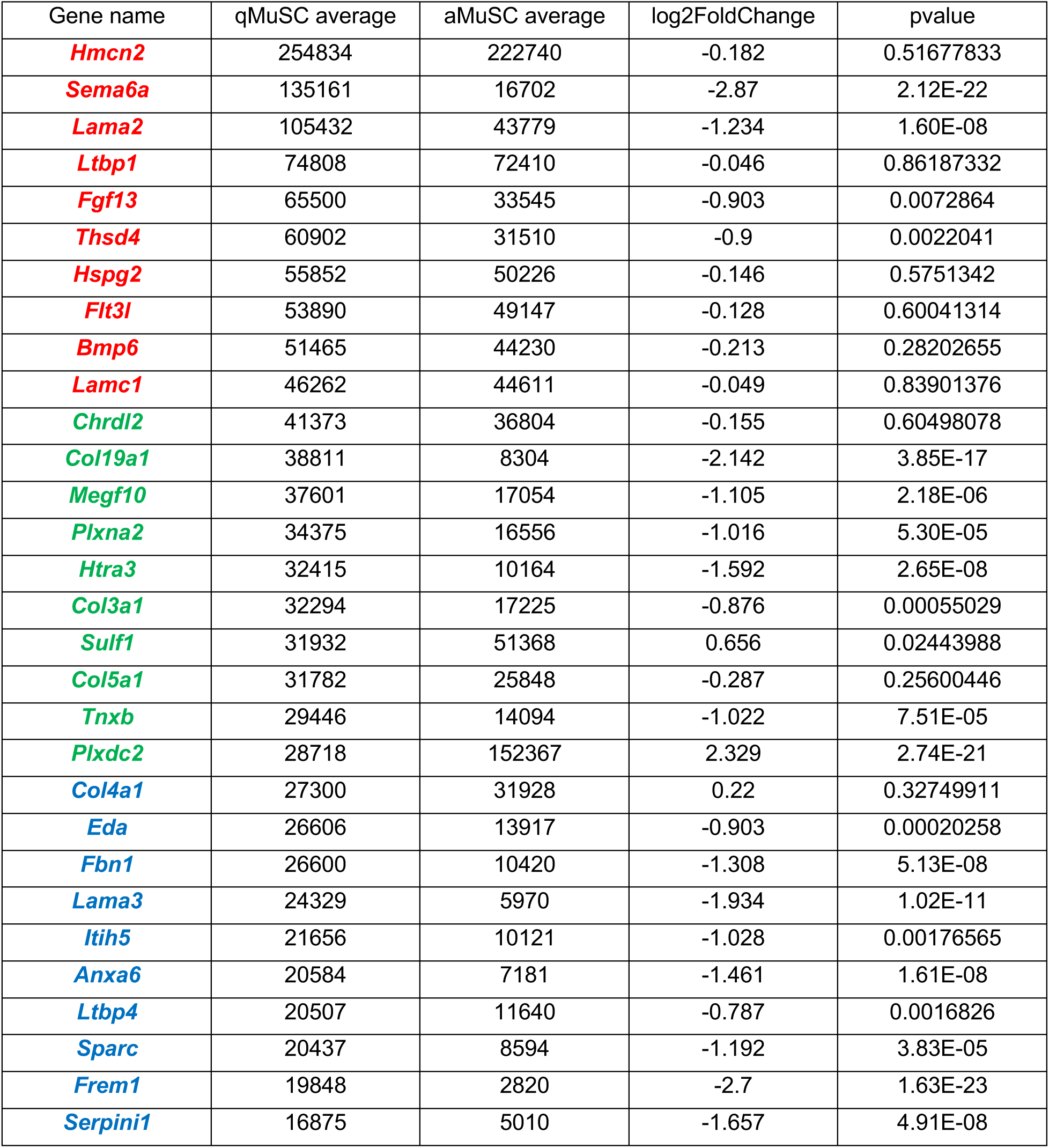
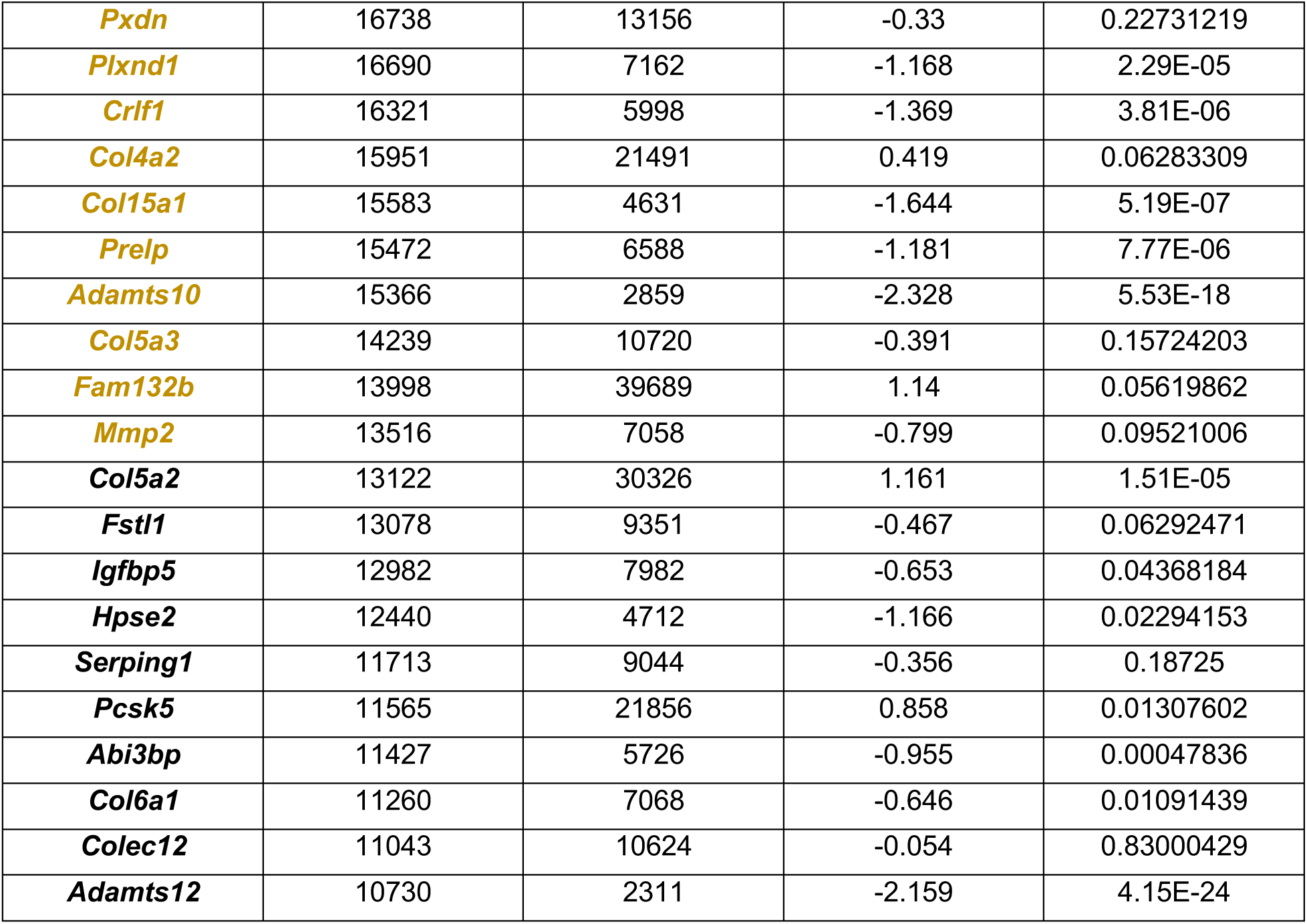
Average expression (number of reads) of the Top 50 matrisome genes expressed by MuSC when quiescent (q MuSC) and upon activation (aMuSC). Log2Foldchange in expression between aMuSC/qMuSC states for these genes and the associated pvalue are also indicated. Gene rank color coding is the same than the one used in Figure 1

| Gene name | qMuSC average | aMuSC average | log2FoldChange | pvalue |
| --- | --- | --- | --- | --- |
| *Hmcn2* | 254834 | 222740 | -0.182 | 0.51677833 |
| *Sema6a* | 135161 | 16702 | -2.87 | 2.12E-22 |
| *Lama2* | 105432 | 43779 | -1.234 | 1.60E-08 |
| *Ltbp1* | 74808 | 72410 | -0.046 | 0.86187332 |
| *Fgf13* | 65500 | 33545 | -0.903 | 0.0072864 |
| *Thsd4* | 60902 | 31510 | -0.9 | 0.0022041 |
| *Hspg2* | 55852 | 50226 | -0.146 | 0.5751342 |
| *Flt3l* | 53890 | 49147 | -0.128 | 0.60041314 |
| *Bmp6* | 51465 | 44230 | -0.213 | 0.28202655 |
| *Lamc1* | 46262 | 44611 | -0.049 | 0.83901376 |
| *Chrdl2* | 41373 | 36804 | -0.155 | 0.60498078 |
| *Col19a1* | 38811 | 8304 | -2.142 | 3.85E-17 |
| *Megf10* | 37601 | 17054 | -1.105 | 2.18E-06 |
| *Plxna2* | 34375 | 16556 | -1.016 | 5.30E-05 |
| *Htra3* | 32415 | 10164 | -1.592 | 2.65E-08 |
| *Col3a1* | 32294 | 17225 | -0.876 | 0.00055029 |
| *Sulf1* | 31932 | 51368 | 0.656 | 0.02443988 |
| *Col5a1* | 31782 | 25848 | -0.287 | 0.25600446 |
| *Tnxb* | 29446 | 14094 | -1.022 | 7.51E-05 |
| *Plxdc2* | 28718 | 152367 | 2.329 | 2.74E-21 |
| *Col4a1* | 27300 | 31928 | 0.22 | 0.32749911 |
| *Eda* | 26606 | 13917 | -0.903 | 0.00020258 |
| *Fbn1* | 26600 | 10420 | -1.308 | 5.13E-08 |
| *Lama3* | 24329 | 5970 | -1.934 | 1.02E-11 |
| *Itih5* | 21656 | 10121 | -1.028 | 0.00176565 |
| *Anxa6* | 20584 | 7181 | -1.461 | 1.61E-08 |
| *Ltbp4* | 20507 | 11640 | -0.787 | 0.0016826 |
| *Sparc* | 20437 | 8594 | -1.192 | 3.83E-05 |
| *Frem1* | 19848 | 2820 | -2.7 | 1.63E-23 |
| *Serpini1* | 16875 | 5010 | -1.657 | 4.91E-08 |
| *Pxdn* | 16738 | 13156 | -0.33 | 0.22731219 |
| *Plxnd1* | 16690 | 7162 | -1.168 | 2.29E-05 |
| *Crlf1* | 16321 | 5998 | -1.369 | 3.81E-06 |
| *Col4a2* | 15951 | 21491 | 0.419 | 0.06283309 |
| *Col15a1* | 15583 | 4631 | -1.644 | 5.19E-07 |
| *Prelp* | 15472 | 6588 | -1.181 | 7.77E-06 |
| *Adamts10* | 15366 | 2859 | -2.328 | 5.53E-18 |
| *Col5a3* | 14239 | 10720 | -0.391 | 0.15724203 |
| *Fam132b* | 13998 | 39689 | 1.14 | 0.05619862 |
| *Mmp2* | 13516 | 7058 | -0.799 | 0.09521006 |
| *Col5a2* | 13122 | 30326 | 1.161 | 1.51E-05 |
| *Fstl1* | 13078 | 9351 | -0.467 | 0.06292471 |
| *Igfbp5* | 12982 | 7982 | -0.653 | 0.04368184 |
| *Hpse2* | 12440 | 4712 | -1.166 | 0.02294153 |
| *Serping1* | 11713 | 9044 | -0.356 | 0.18725 |
| *Pcsk5* | 11565 | 21856 | 0.858 | 0.01307602 |
| *Abi3bp* | 11427 | 5726 | -0.955 | 0.00047836 |
| *Col6a1* | 11260 | 7068 | -0.646 | 0.01091439 |
| *Colec12* | 11043 | 10624 | -0.054 | 0.83000429 |
| *Adamts12* | 10730 | 2311 | -2.159 | 4.15E-24 |

**Table S2 related to Figure 5.**
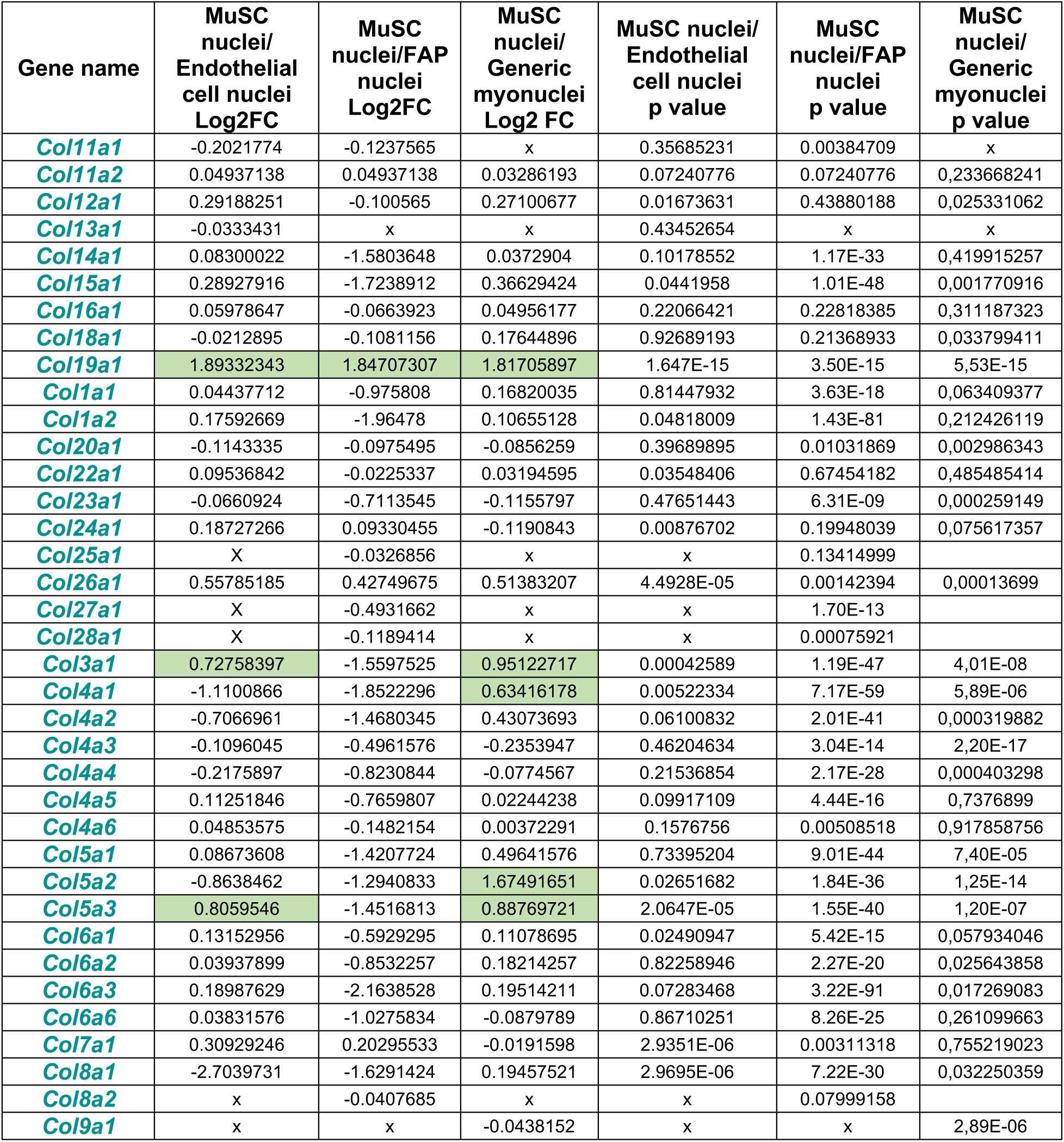
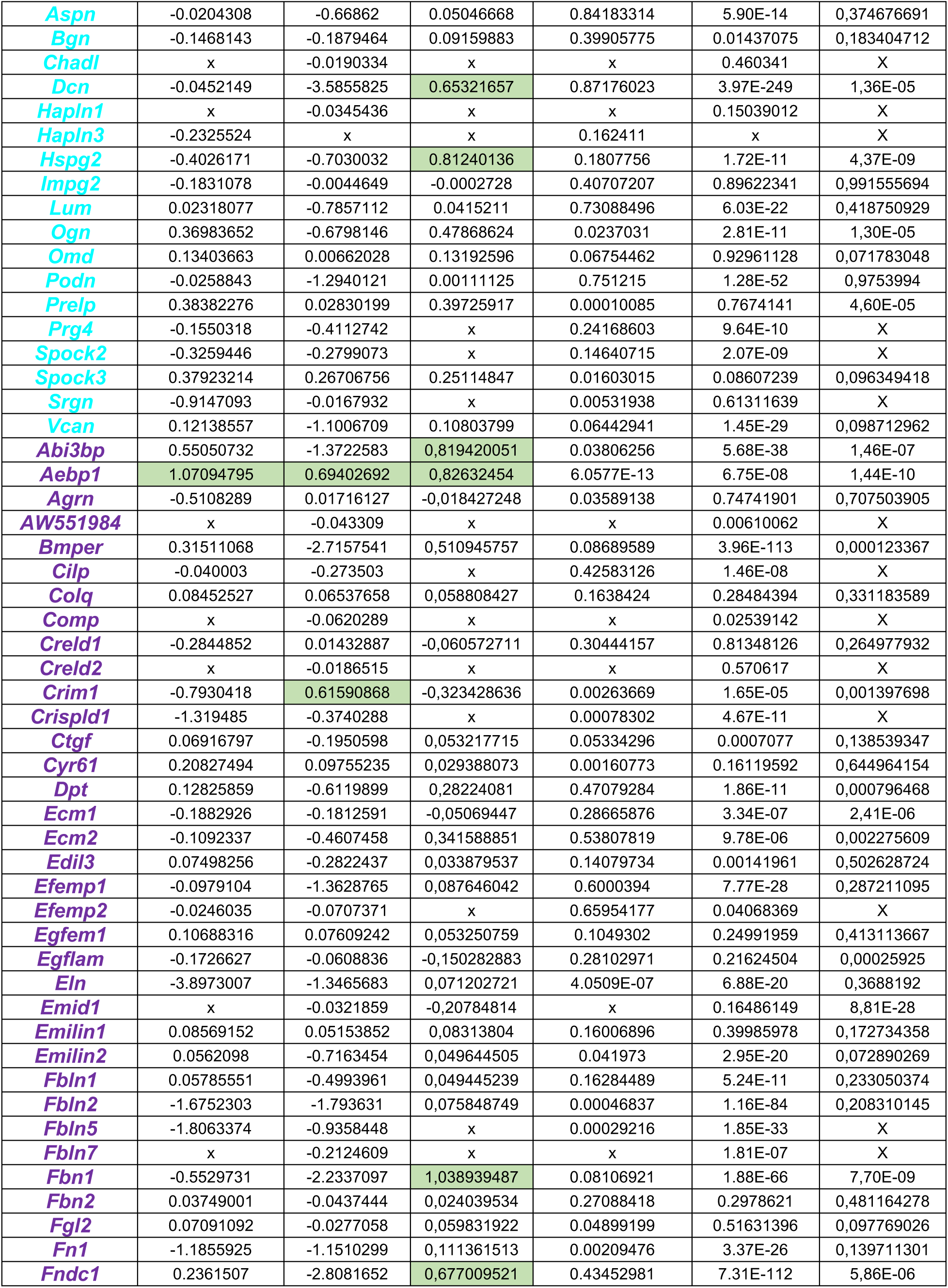
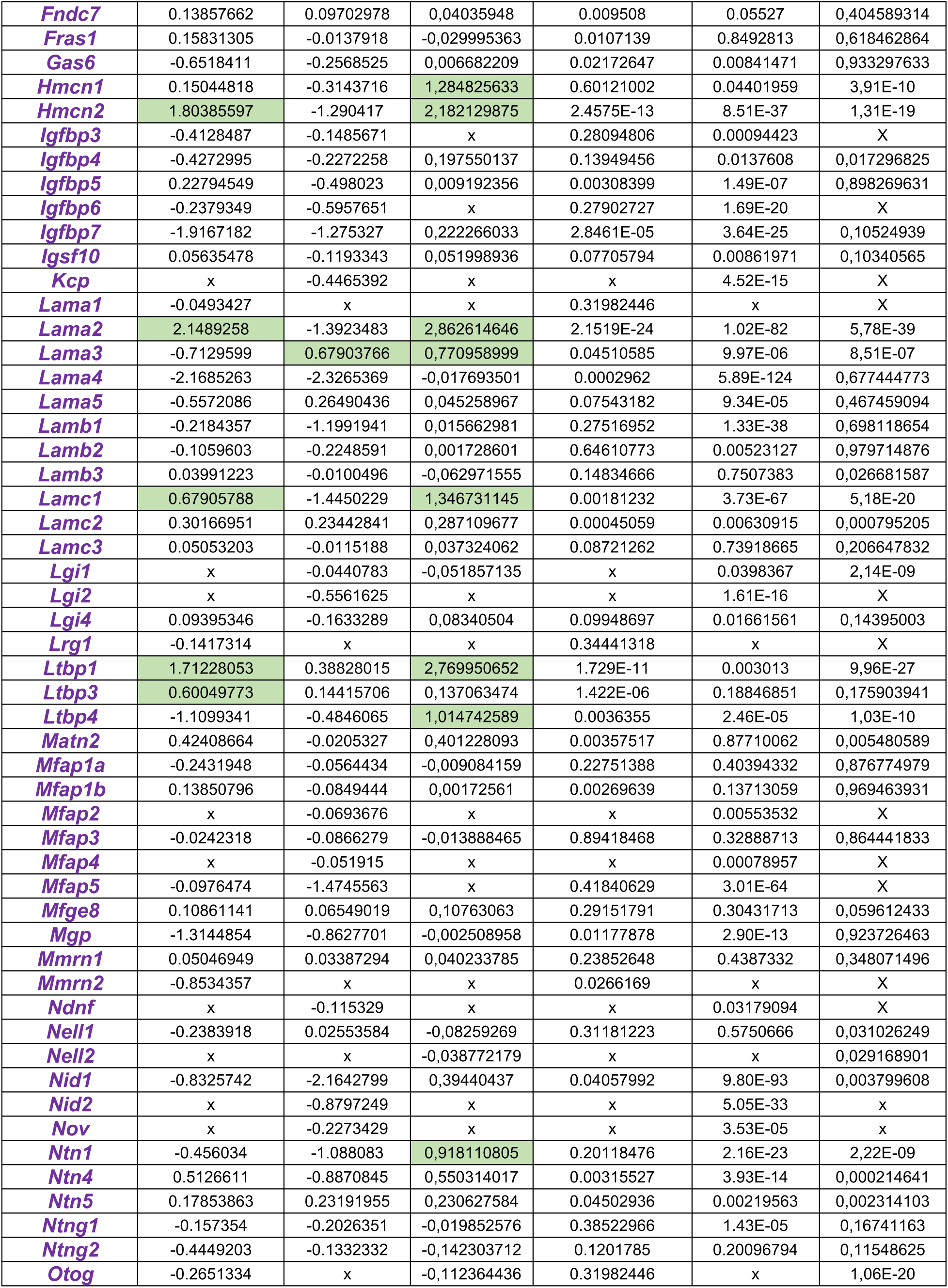
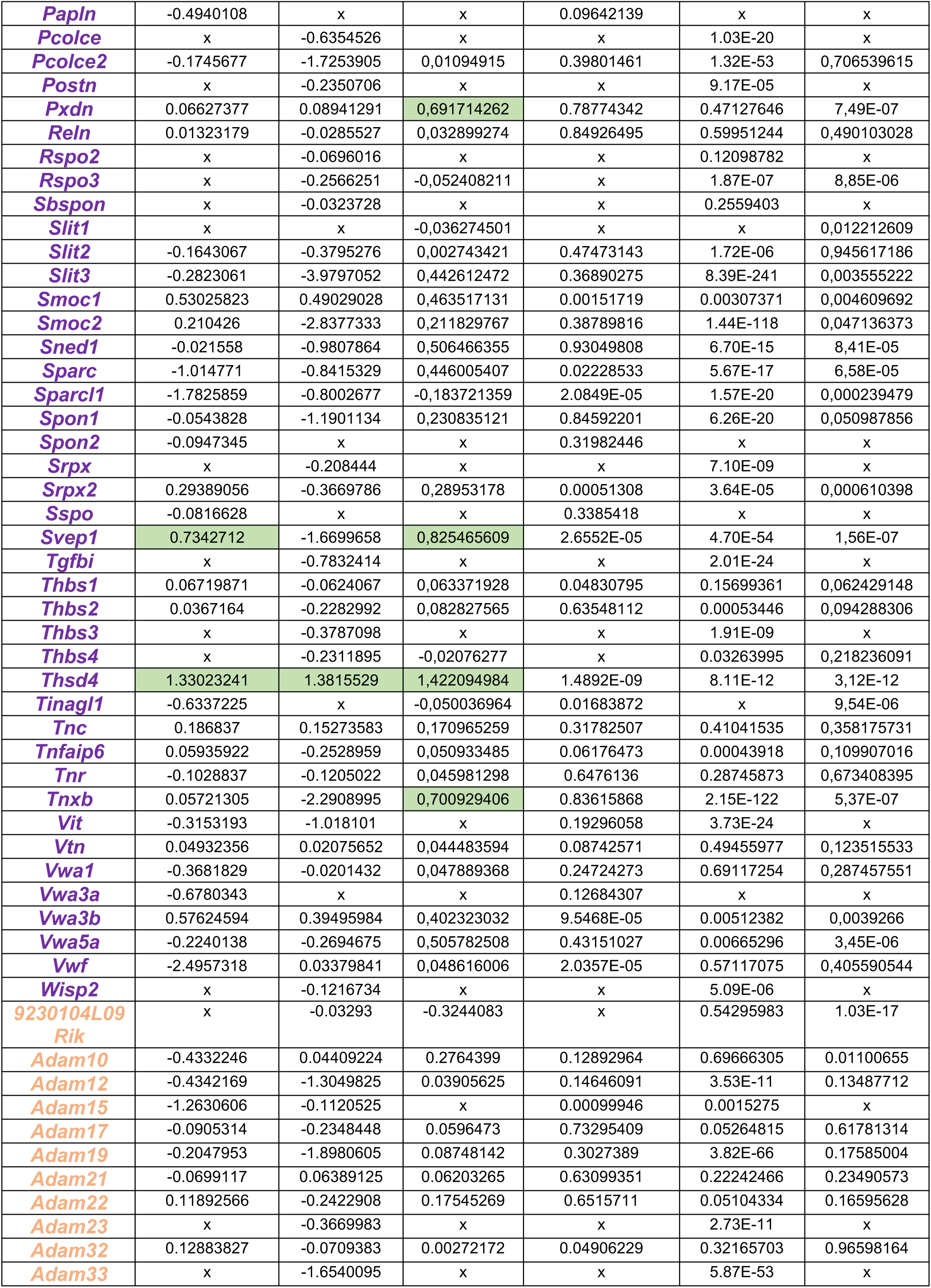
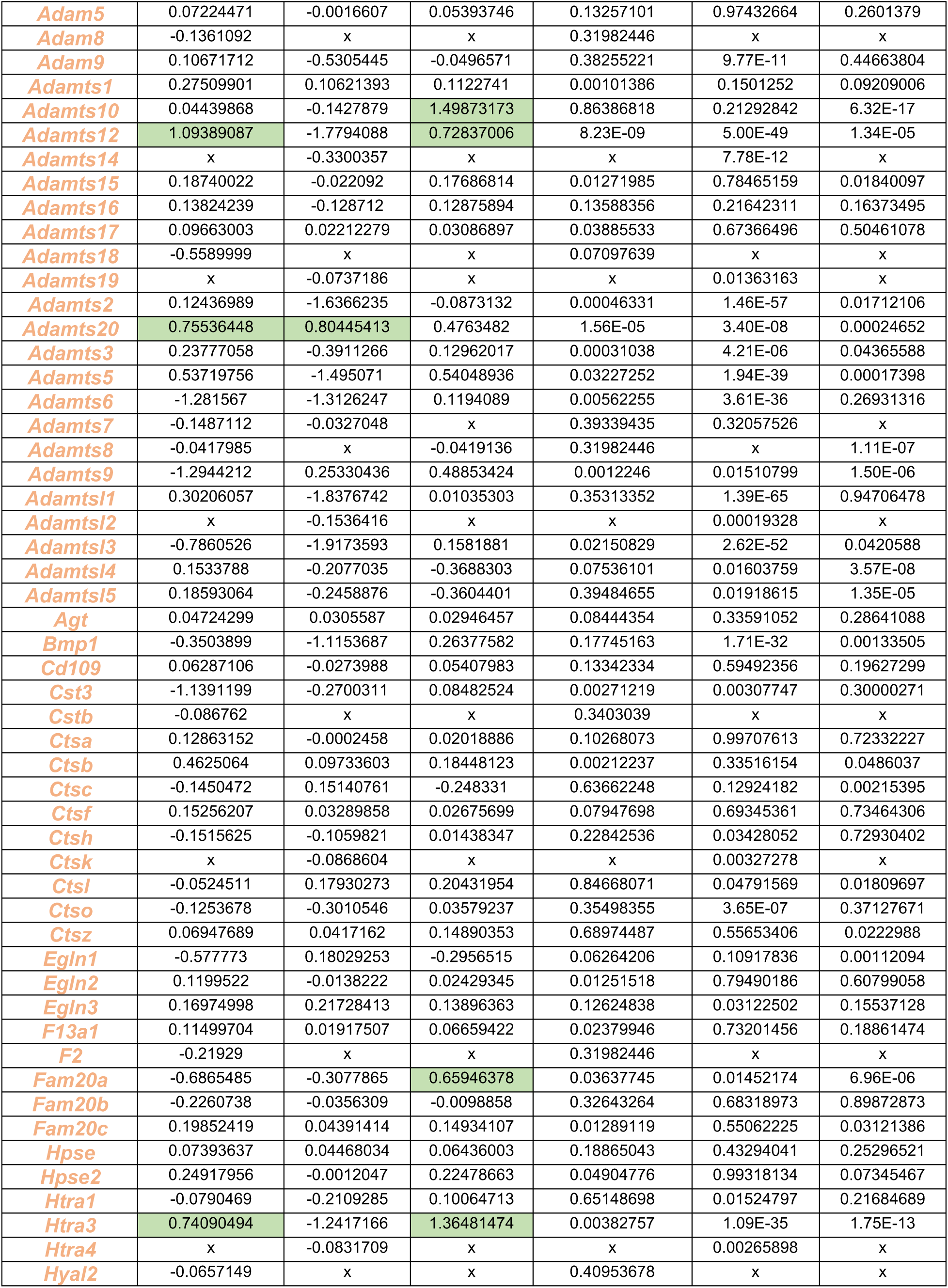
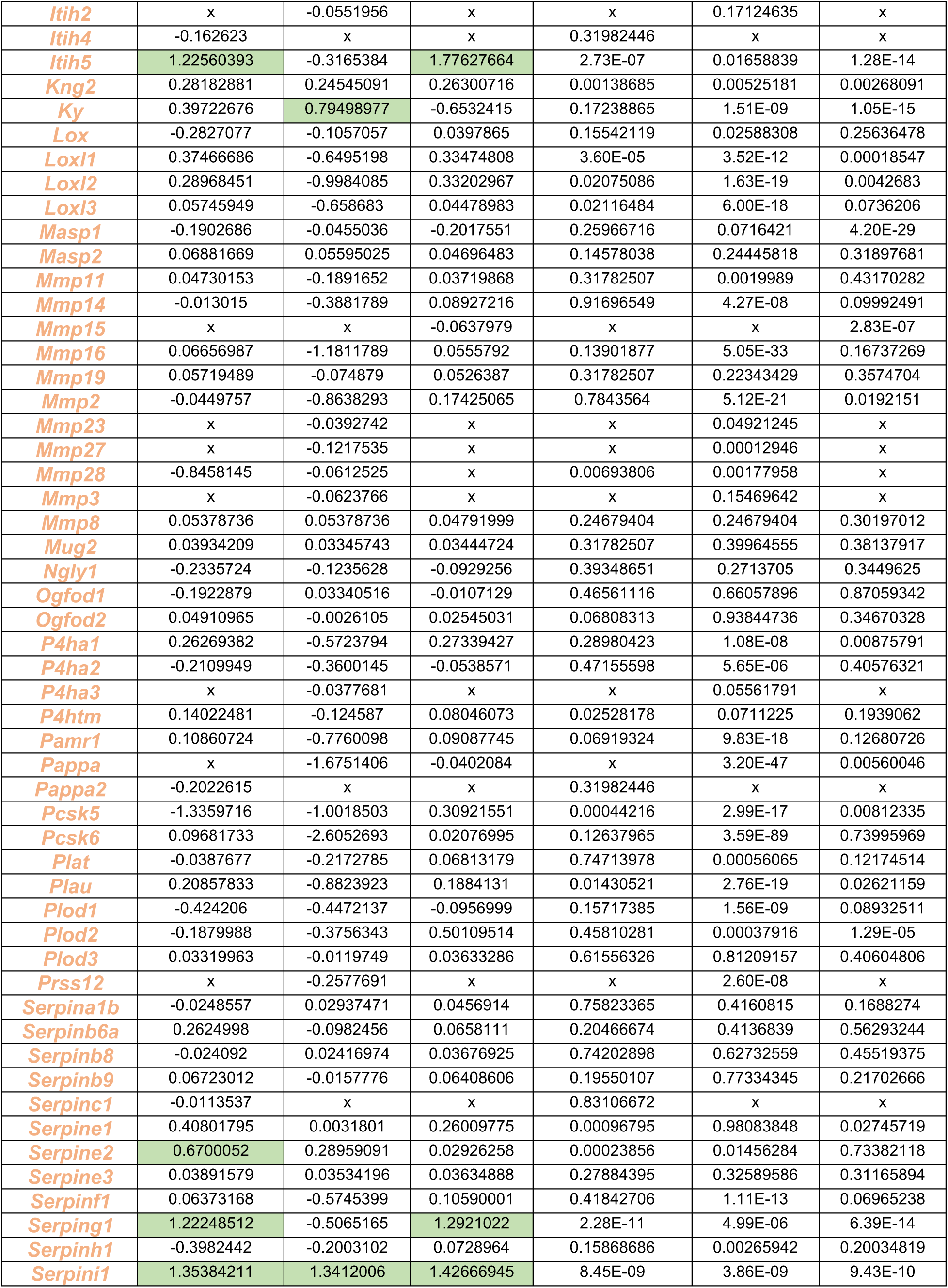
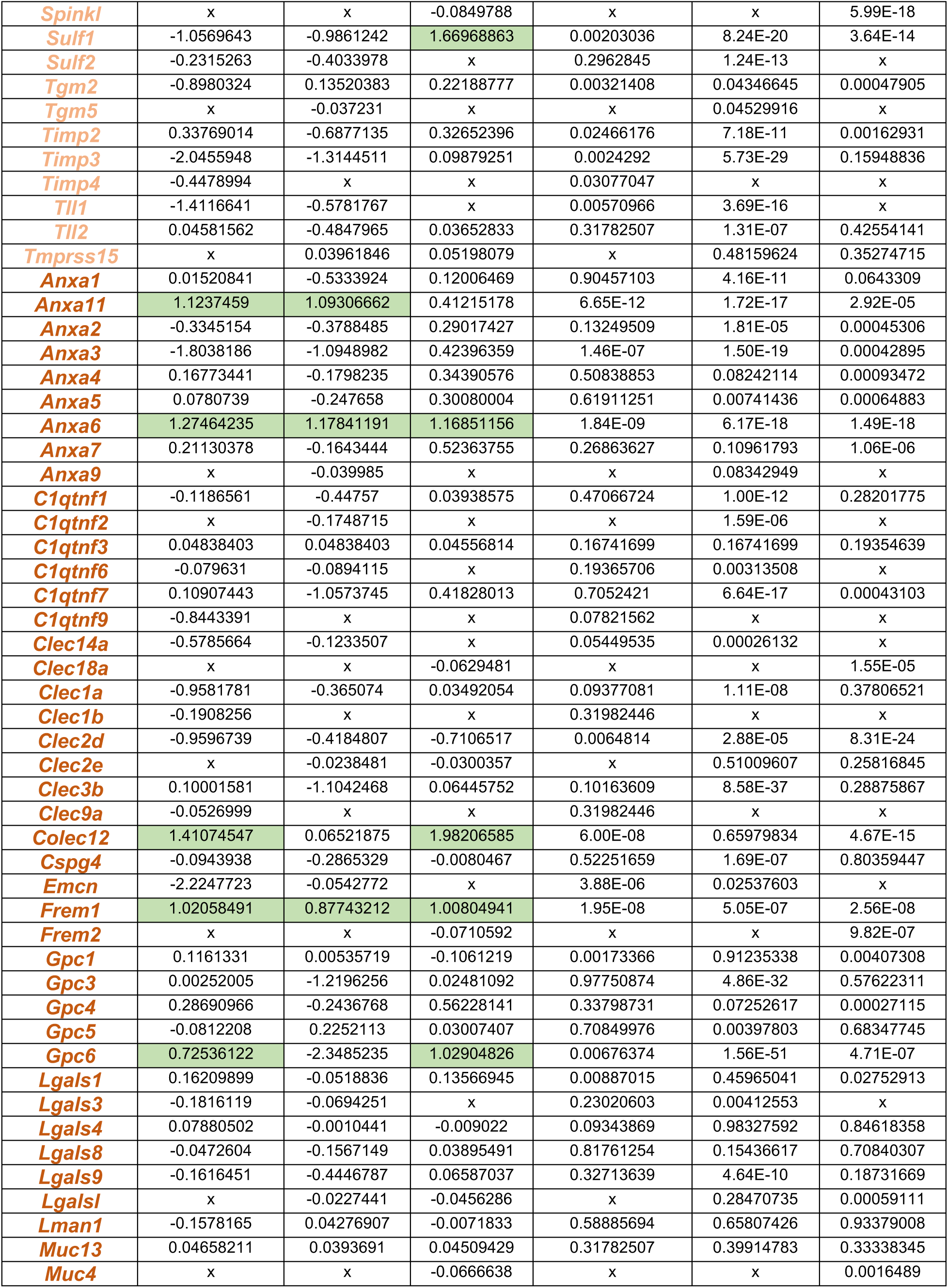
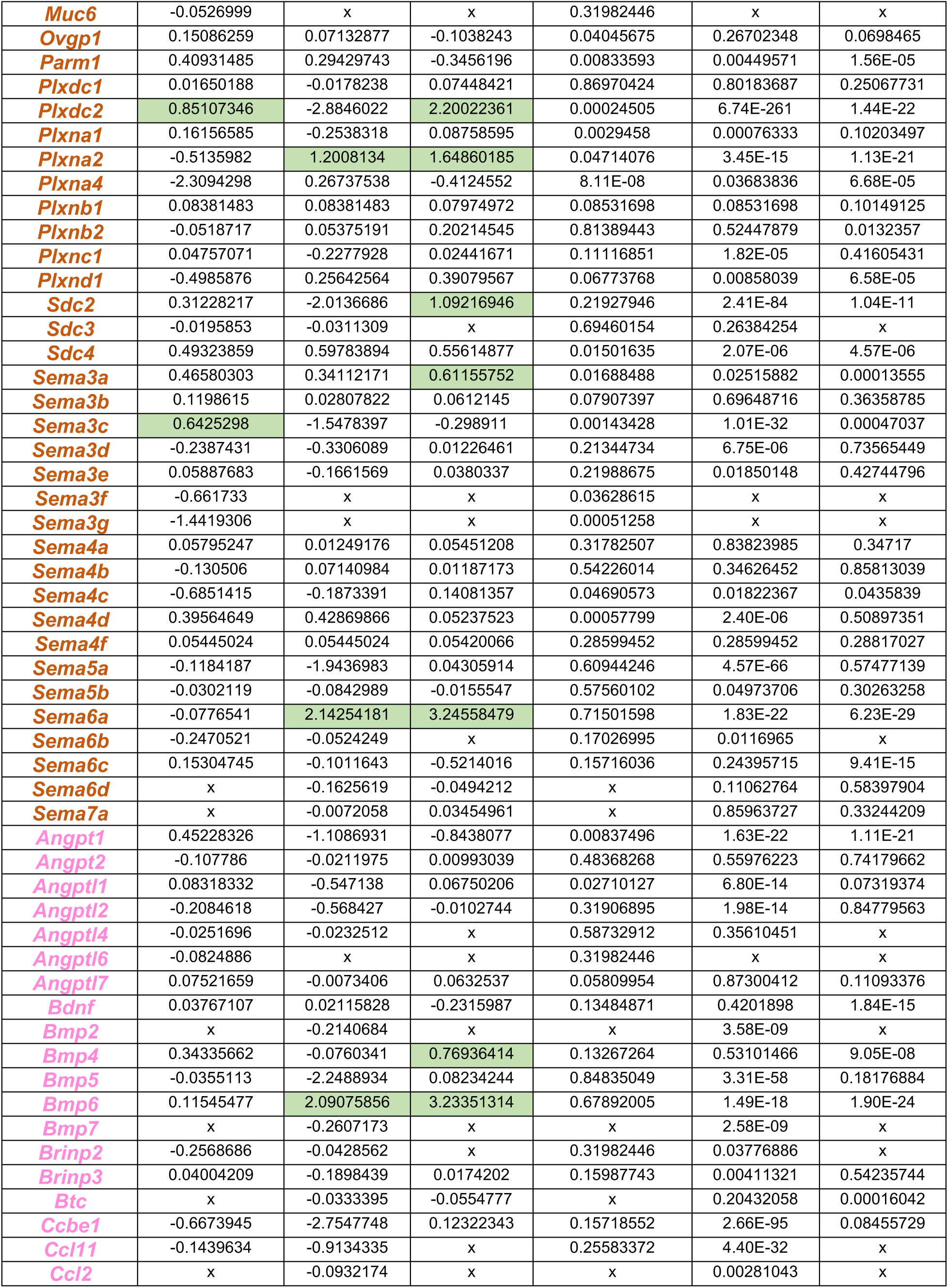
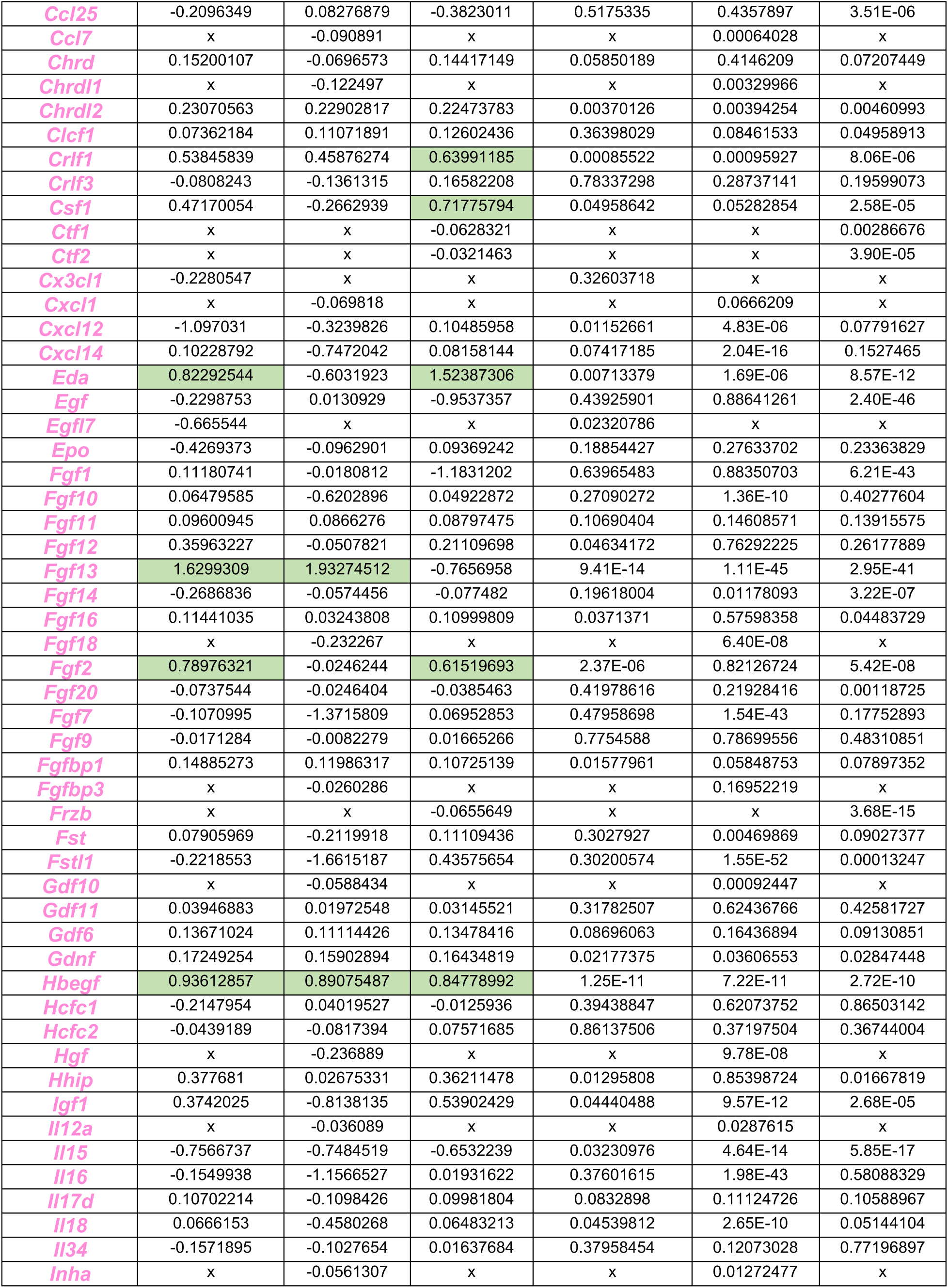
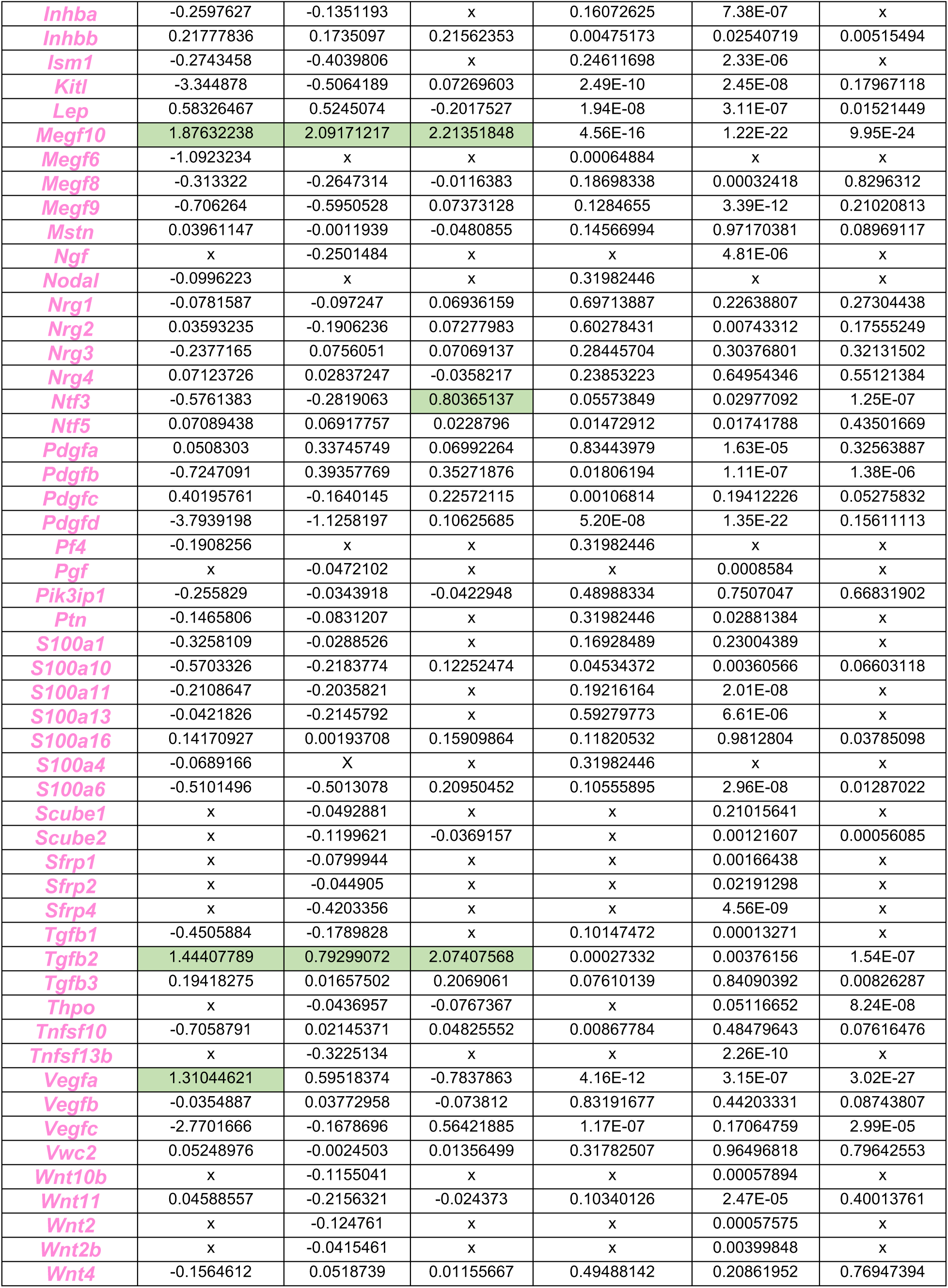
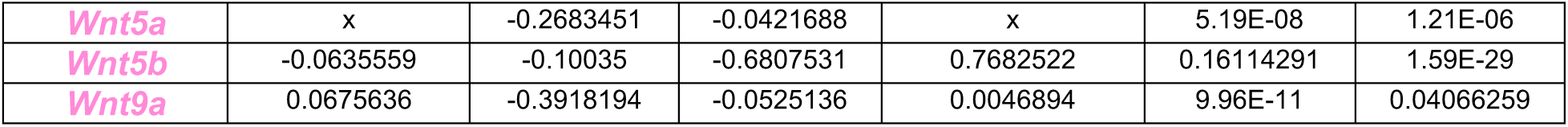
Pairwise comparison of matrisome genes expression levels between MuSC and a given neighboring cell (FAP, Myofiber, Endothelial cell). Analysis performed on the single nuclei RNA seq dataset published in (Machado et al., 2021.) Green cell color marks a log2FC ≥ 0.6 and therefore a significant enrichment in gene expression level in MuSC compared to the given neighboring cells. A gene for which the 3 first cells are green is therefore considered as a MuSC matrisome signature gene. x = comparison was not possible as the average expression level was under the set expression threshold (0.026) for both cell types. The gene color coding indicating matrisome categories is the same than in Figure 1 A.

**Table S3 related to Fig 7.**
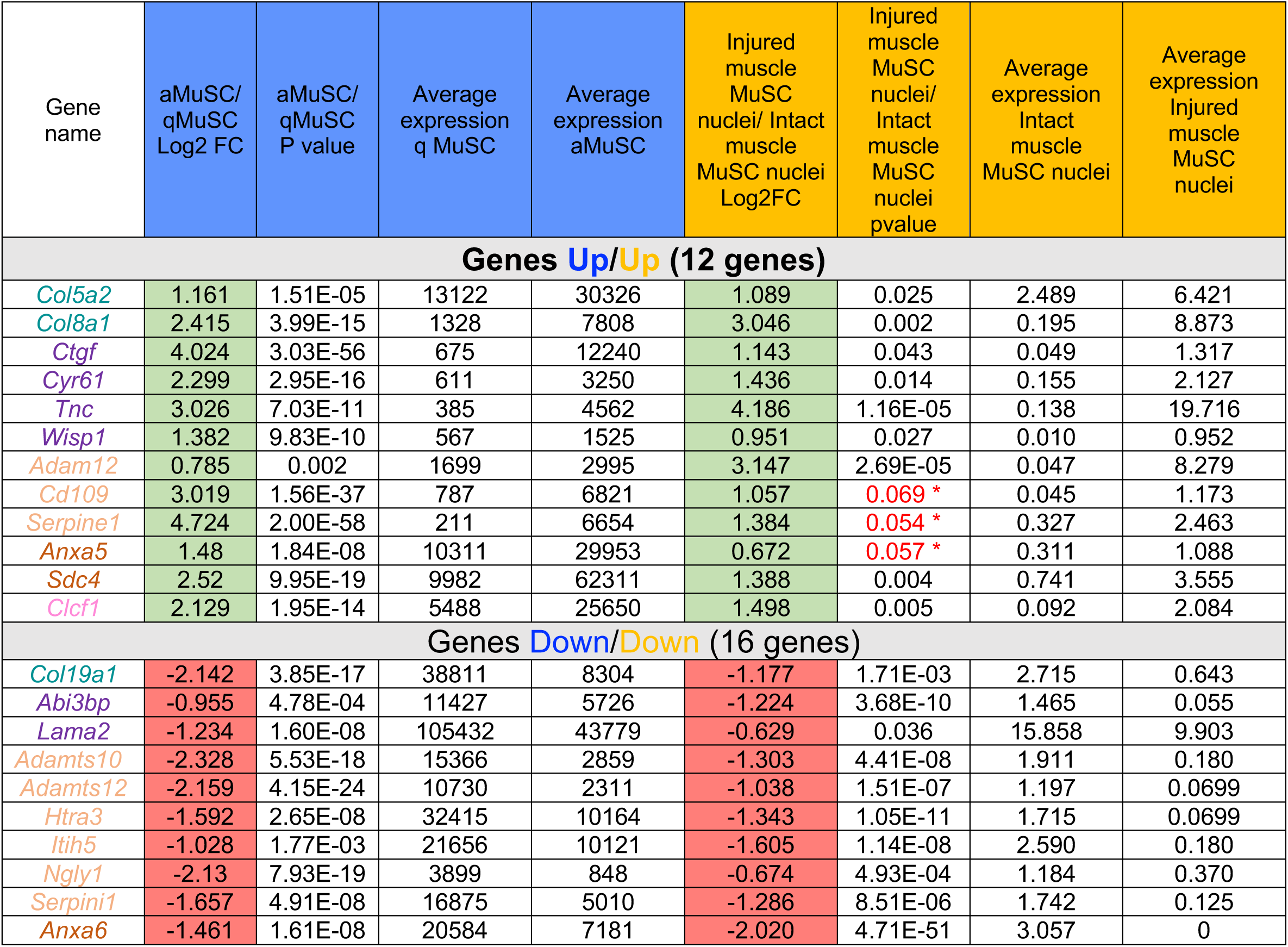
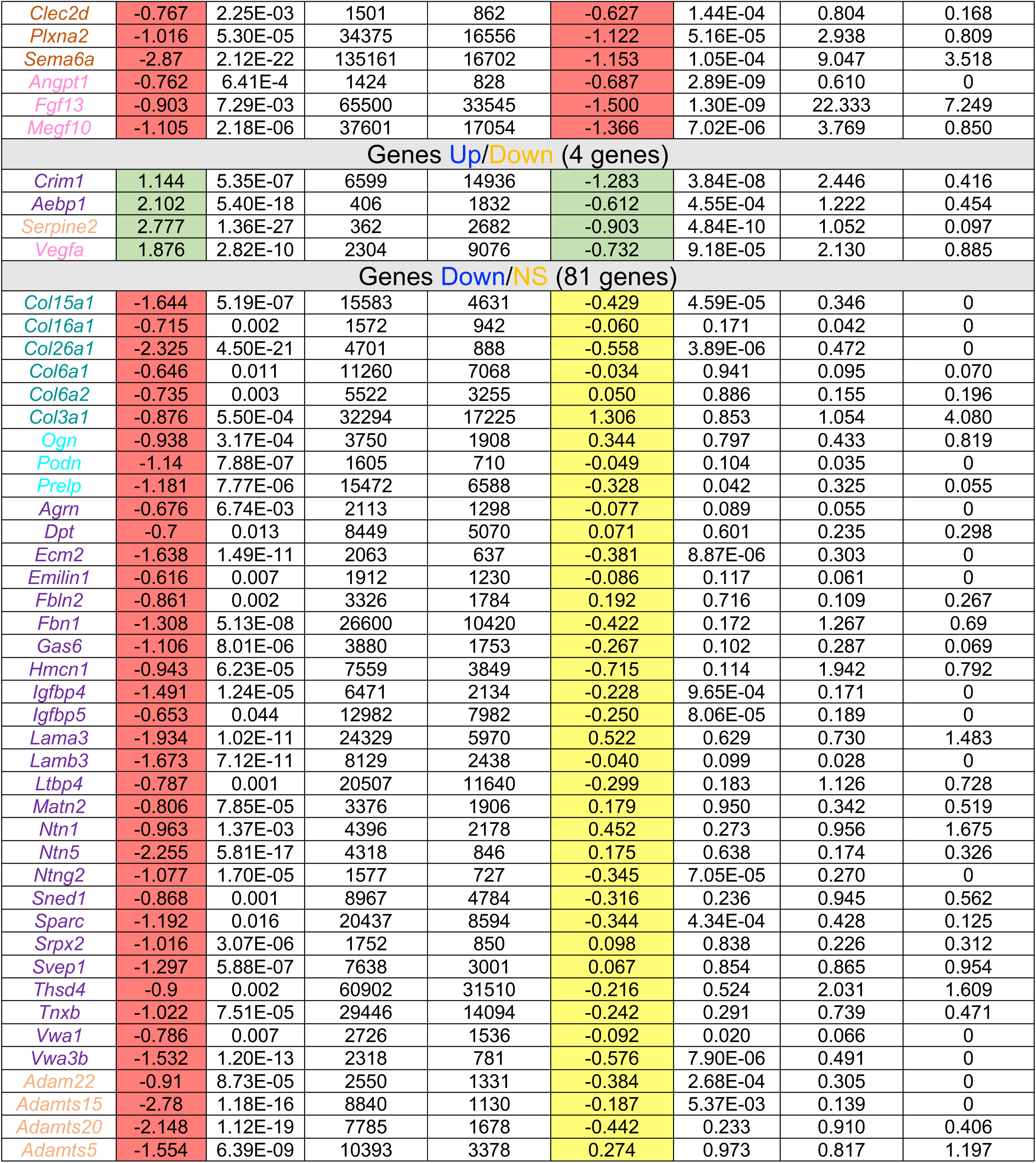
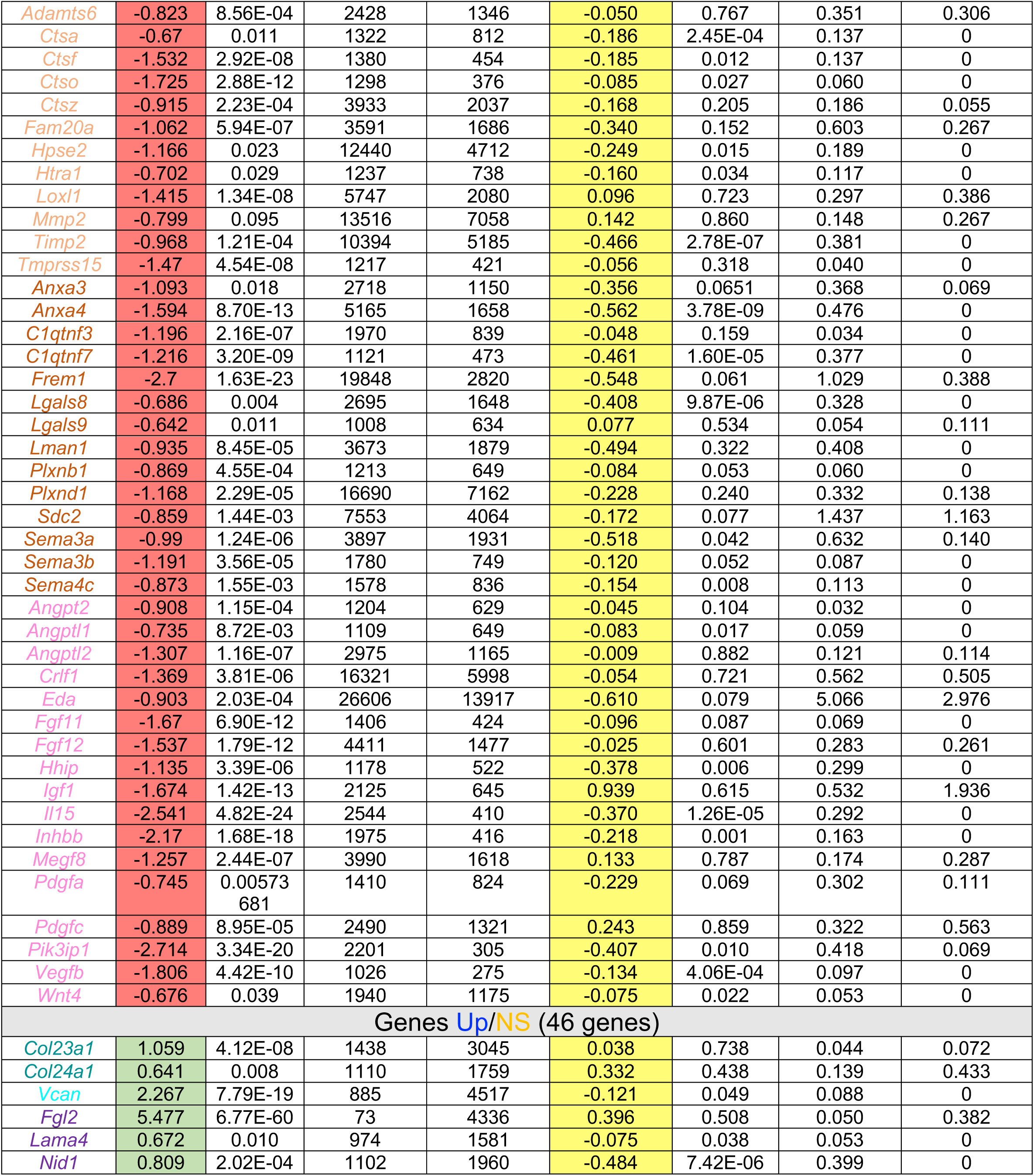
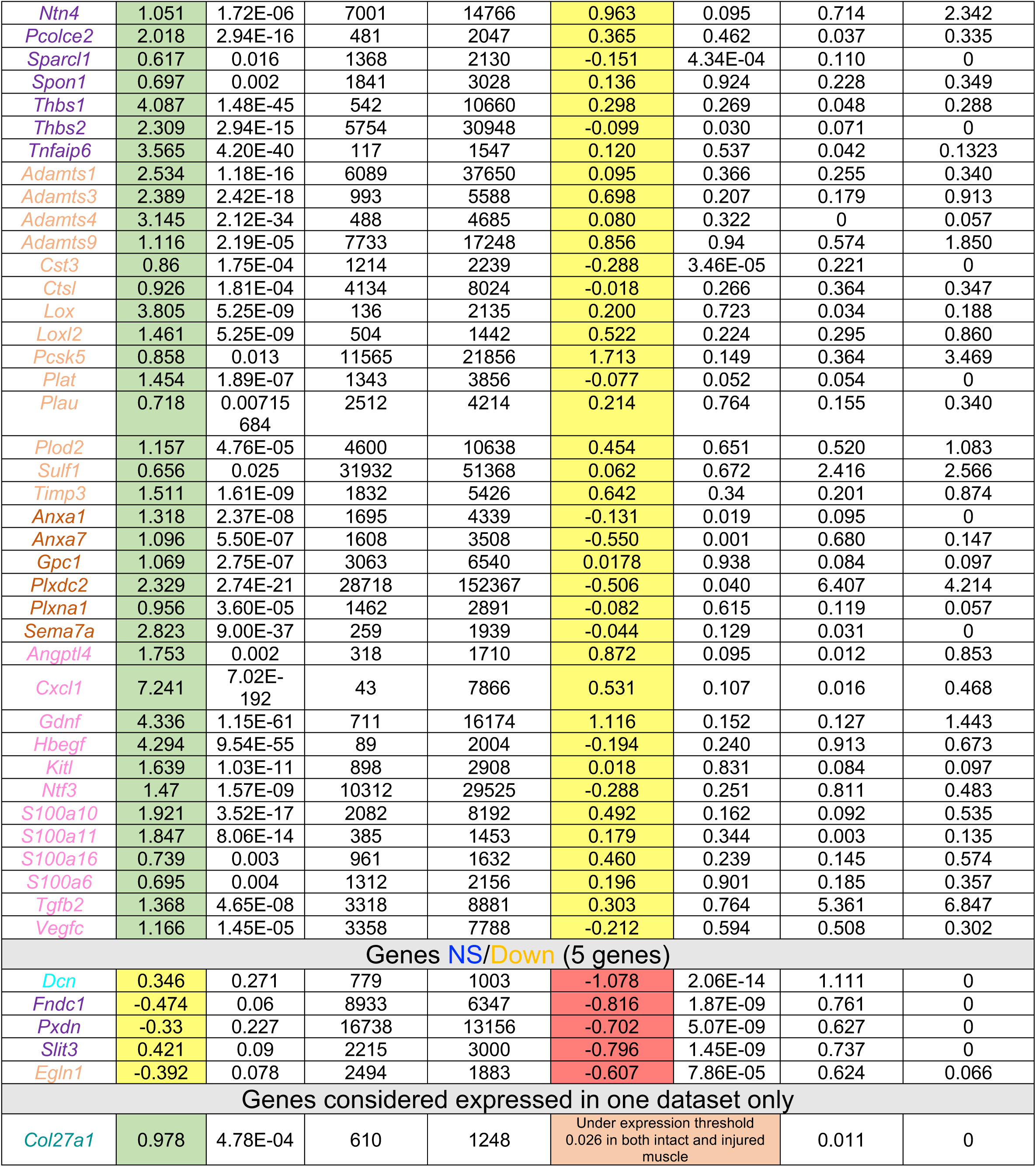
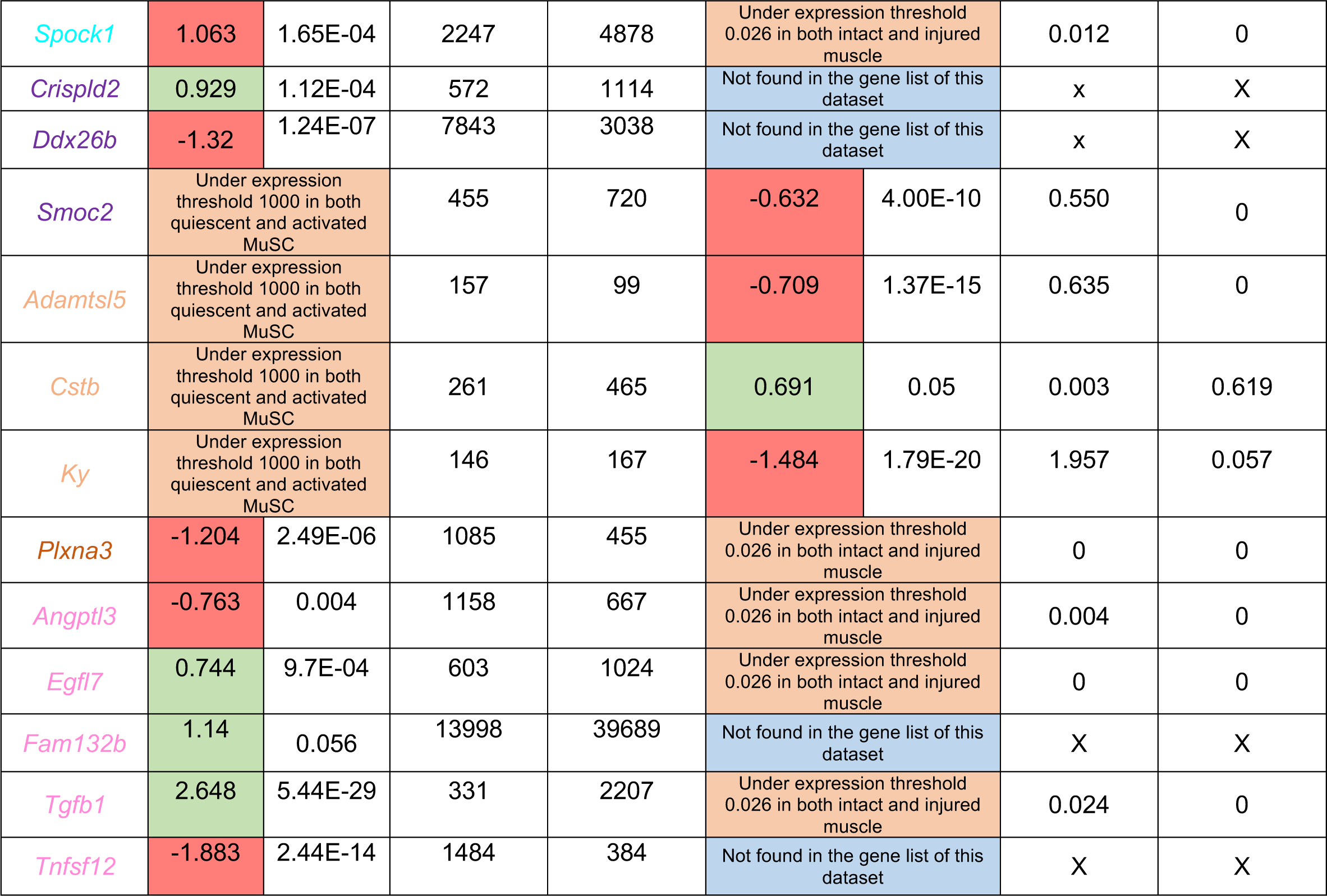
Comparison of Matrisome genes regulation upon MuSC activation during tissue dissociation and tissue injury from publicly available data sets (Machado et al., 2021). The blue columns indicate the Log2Foldchange in gene expression between aMuSC/qMuSC states, the associated pvalue, the average gene expression (number of reads) by qMuSC and aMuSC. Only the matrisome genes which are significantly regulated in at least one situation are shown. Genes were classified into 7 categories depending on their relative regulation status in tissue dissociation and tissue injury: Genes up in both cases; Genes down in both cases; Genes inversely regulated; Genes upregulated or downregulated in one case but not significantly regulated in the other; Genes dysregulated in one dataset but considered as not being expressed by MuSC in the other. Green, significantly upregulated genes (log2FC ≥ 0.6 and p value <0.07). Red, significantly downregulated genes (log2FC < −0.6 and pvalue<0.07). Yellow, not differentially expressed genes (|log2 FC| < 0.6 or pvalue > 0.07). Stars indicate 0.05 < p value < 0.07. The gene color coding indicating matrisome categories is the same than in Figure 1 A.

## REFERENCES

Baghdadi M.B., D. Castel, L. Machado, S. Fukada, D. E. Birk, F. Relaix, S. Tajbakhsh, and P. Mourikis. 2018. Reciprocal Signalling by Notch–Collagen V–CALCR Retains Muscle Stem Cells in Their Niche. Nature 557:714–18. 10.1038/s41586-018-0144-9.

Benjamini Y., and Y. Hochberg. 1995. Controlling the False Discovery Rate: A Practical and Powerful Approach to Multiple Testing. Journal of the Royal Statistical Society: Series B (Methodological) 57: 89–300. 10.1111/j.2517-6161.1995.tb02031.x.

Bizzarro V., Belvedere, R., Dal Piaz, F., Parente, L., Petrella, A., 2012. Annexin A1 Induces Skeletal Muscle Cell Migration Acting through Formyl Peptide Receptors. PLoS ONE 7, e48246. 10.1371/journal.pone.0048246

Blackburn P.R., Z. Xu, K.E. Tumelty, R.W. Zhao, W.J. Monis, K.G. Harris, J.M. Gass, et al. 2018. Bi-Allelic Alterations in AEBP1 Lead to Defective Collagen Assembly and Connective Tissue Structure Resulting in a Variant of Ehlers-Danlos Syndrome.The American Journal of Human Genetics 102: 696–705. 10.1016/j.ajhg.2018.02.018.

Borges V.M., T. W. Lee, D. L. Christie and N. P. Birch. 2010. Neuroserpin regulates the density of dendritic protrusions and dendritic spine shape in cultured hippocampal neurons. J. Neurosci. Res. 88: 2610–2617. 10.1111/j.1471-4159.2012.07655.x

Bonod-Bidaud C., M. Roulet, U. Hansen, A. Elsheikh, M. Malbouyres, S. Ricard-Blum, C. Faye, E. Vaganay, P. Rousselle, F. Ruggiero. 2012. In vivo evidence for a bridging role of a collagen V subtype at the epidermis-dermis interface, J. Invest. Dermatol. 132 :1841– 1849. doi:10.1038/jid.2012.56.

Calvo A.C., L. Moreno, L. Moreno, J.M. Toivonen, R.l Manzano, N. Molina, M. de la Torre, et al. 2020. Type XIX Collagen: A Promising Biomarker from the Basement Membranes. Neural Regeneration Research 15: 988. 10.4103/1673-5374.270299.

Chen L-H., C-Y. Liao, L-C. Lai, M-H. Tsai, and E.Y. Chuang. 2019. Semaphorin 6A Attenuates the Migration Capability of Lung Cancer Cells via the NRF2/HMOX1 Axis. Scientific Reports 9:13302. 10.1038/s41598-019-49874-8.

Cheng T-W., and Q. Gong. 2009. Secreted TARSH Regulates Olfactory Mitral Cell Dendritic Complexity. European Journal of Neuroscience 2:1083–95. 10.1111/j.1460-9568.2009.06660.x.

Cheng Z., Y. Zhang, Y. Tian, Y. Chen, F. Ding, H. Wu, Y. Ji, and M. Shen. 2021. Cyr61 Promotes Schwann Cell Proliferation and Migration via Αvβ3 Integrin. BMC Molecular and Cell Biology 22: 21. 10.1186/s12860-021-00360-y.

Clementz A.G., and A. Harris. 2013. Collagen XV: Exploring Its Structure and Role within the Tumor Microenvironment. Molecular Cancer Research 11:1481–86. 10.1158/1541-7786.MCR-12-0662.

Crisponi L., I. Buers, and F. Rutsch. 2022. CRLF1 and CLCF1 in Development, Health and Disease. International Journal of Molecular Sciences 23:992. 10.3390/ijms23020992.

Csapo M. Gumpenberger, and B. Wessner. 2020. Skeletal Muscle Extracellular Matrix – What Do We Know About Its Composition, Regulation, and Physiological Roles? A Narrative Review. Frontiers in Physiology 11: 253. 10.3389/fphys.2020.00253.

Daniels M.P. 2012. The Role of Agrin in Synaptic Development, Plasticity and Signaling in the Central Nervous System. Neurochemistry International 61:848–53. 10.1016/j.neuint.2012.02.028.

De Micheli A.J., E.J. Laurilliard, C.L. Heinke, H. Ravichandran, P. Fraczek, S. Soueid-Baumgarten, I. De Vlaminck, O. Elemento, and B.D. Cosgrove. 2020. Single-Cell Analysis of the Muscle Stem Cell Hierarchy Identifies Heterotypic Communication Signals Involved in Skeletal Muscle Regeneration. Cell Reports 30:3583–3595.e5. 10.1016/j.celrep.2020.02.067.

Flynn L.A., A.R. Blissett, E.P. Calomeni, and G. Agarwal. 2010. Inhibition of Collagen Fibrillogenesis by Cells Expressing Soluble Extracellular Domains of DDR1 and DDR2. Journal of Molecular Biology 395: 533–43. 10.1016/j.jmb.2009.10.073.

Garbe JH, Göhring W, Mann K, Timpl R, Sasaki T. 2002. Complete sequence, recombinant analysis and binding to laminins and sulphated ligands of the N-terminal domains of laminin α3B and α5 chains. Biochemical Journal 362: 213–221.10.1042/bj3620213

Iversen K., F. Beaubien, J.E.A. Prince, and J-F. Cloutier. 2020. Axon Guidance: Slit–Robo Signaling. In Cellular Migration and Formation of Axons and Dendrites, 147–73. Elsevier. 10.1016/B978-0-12-814407-7.00007-9.

Jones C.P., S.C. Pitchford, C.M. Lloyd, and S.M. Rankin. 2009. CXCR2 Mediates the Recruitment of Endothelial Progenitor Cells During Allergic Airways Remodeling. Stem Cells 27: 3074–81. 10.1002/stem.222.

Kann A.P., Hung, M., Wang, W., Nguyen, J., Gilbert, P.M., Wu, Z., Krauss, R.S., 2022. An injury-responsive Rac-to-Rho GTPase switch drives activation of muscle stem cells through rapid cytoskeletal remodeling. Cell Stem Cell 29:933–947.e6. 10.1016/j.stem.2022.04.016

Keeley D.P. and D.R. Sherwood. 2019. Tissue Linkage through Adjoining Basement Membranes: The Long and the Short Term of It. Matrix Biology 75–76: 58–71. 10.1016/j.matbio.2018.05.009.

Kroeze K.L., M.A. Boink, S.C. Sampat-Sardjoepersad, T. Waaijman, R.J. Scheper, and S. Gibbs. 2012. Autocrine Regulation of Re-Epithelialization After Wounding by Chemokine Receptors CCR1, CCR10, CXCR1, CXCR2, and CXCR3. Journal of Investigative Dermatology 132: 216–25. 10.1038/jid.2011.245.

Larouche J.A., M. Mohiuddin, J.J. Choi, P.J. Ulintz, P. Fraczek, K. Sabin, S. Pitchiaya, et al. 2021. Murine Muscle Stem Cell Response to Perturbations of the Neuromuscular Junction Are Attenuated with Aging. ELife 10:e66749. 10.7554/eLife.66749.

Lepper C, T.A. Partridge, C-M Fan 2011. An absolute requirement for Pax7-positive satellite cells in acute injury-induced skeletal muscle regeneration. Development 138:3639–46. doi: 10.1242/dev.067595.

Liu W., A. Klose, S. Forman, N.D. Paris, L Wei-LaPierre, M. Cortés-Lopéz, A. Tan, et al. 2017. Loss of Adult Skeletal Muscle Stem Cells Drives Age-Related Neuromuscular Junction Degeneration. ELife 6: e26464. 10.7554/eLife.26464.

Liu W., L. Wei-LaPierre, A. Klose, R.T. Dirksen, and J.V. Chakkalakal. 2015. Inducible Depletion of Adult Skeletal Muscle Stem Cells Impairs the Regeneration of Neuromuscular Junctions. ELife 4: e09221. 10.7554/eLife.09221.

Love, M.I., W. Huber, and S. Anders. 2014. Moderated Estimation of Fold Change and Dispersion for RNA-Seq Data with DESeq2. Genome Biology 15:550. 10.1186/s13059-014-0550-8.

Ma N.C., D. Chen, J.-H. Lee,, P. Kuri, E.B. Hernandez,, J. Kocan, H. Mahmood, E.D. Tichy, P Rompolas, F. Mourkioti. 2022. Piezo1 regulates the regenerative capacity of skeletal muscles via orchestration of stem cell morphological states. Science Advances 8 :eabn0485. 10.1126/sciadv.abn0485

Machado L., J.E. de Lima, O. Fabre, C. Proux, R. Legendre, A. Szegedi, H. Varet, et al. 2017. In Situ Fixation Redefines Quiescence and Early Activation of Skeletal Muscle Stem Cells. Cell Reports 21:1982–93. 10.1016/j.celrep.2017.10.080.

Machado L., P. Geara, J. Camps, M. Dos Santos, F. Teixeira-Clerc, J. Van Herck, H. Varet, et al. 2021. Tissue Damage Induces a Conserved Stress Response That Initiates Quiescent Muscle Stem Cell Activation. Cell Stem Cell 28:1125–1135.e7. 10.1016/j.stem.2021.01.017.

Maiese K. 2015. Novel Applications of Trophic Factors, Wnt and WISP for Neuronal Repair and Regeneration in Metabolic Disease. Neural Regeneration Research 10:518. 10.4103/1673-5374.155427.

Mehta V., K-L. Pang, D. Rozbesky, K. Nather, A. Keen, D. Lachowski, Y. Kong, et al. 2020. The Guidance Receptor Plexin D1 Is a Mechanosensor in Endothelial Cells. Nature 578:290–95. 10.1038/s41586-020-1979-4.

Mihai C., D.F. Iscru, L.J. Druhan, T.S. Elton, and G. Agarwal. 2006. Discoidin Domain Receptor 2 Inhibits Fibrillogenesis of Collagen Type 1. Journal of Molecular Biology 361:864–76. 10.1016/j.jmb.2006.06.067.

Morote-Garcia J.C., D. Napiwotzky, D. Kohler and P. Rosenberger. 2012. Endothelial Semaphorin 7A Promotes Neutrophil Migration during Hypoxia. Proceedings of the National Academy of Sciences 109:14146–51. 10.1073/pnas.1202165109.

Morrissey M.A., R. Jayadev, G.R. Miley, C.A. Blebea, Q. Chi, S. Ihara, and D.R. Sherwood. 2016. SPARC Promotes Cell Invasion In Vivo by Decreasing Type IV Collagen Levels in the Basement Membrane. PLOS Genetics 12:e1005905. 10.1371/journal.pgen.1005905.

Murach K.A, I.J. Vechetti, D.W. Van Pelt, S.E. Crow, C.M. Dungan, V.C. Figueiredo, K. Kosmac, et al. 2020. Fusion-Independent Satellite Cell Communication to Muscle Fibers During Load-Induced Hypertrophy. Function 1:zqaa009. 10.1093/function/zqaa009.

Naba A., K.R. Clauser, S. Hoersch, H. Liu, S.A. Carr, and R.O. Hynes. 2012. The Matrisome: In Silico Definition and In Vivo Characterization by Proteomics of Normal and Tumor Extracellular Matrices. Molecular & Cellular Proteomics 11:M111.014647. 10.1074/mcp.M111.014647.

Pavlakis E., R. Chiotaki, and G. Chalepakis. 2011. The Role of Fras1/Frem Proteins in the Structure and Function of Basement Membrane. The International Journal of Biochemistry & Cell Biology 43:487–95. 10.1016/j.biocel.2010.12.016.

Rayagiri S.S., D. Ranaldi, A. Raven, N.I. F.M. Azhar, O. Lefebvre, P.S. Zammit, and A-G. Borycki. 2018. Basal Lamina Remodeling at the Skeletal Muscle Stem Cell Niche Mediates Stem Cell Self-Renewal. Nature Communications 9:1075. 10.1038/s41467-018-03425-3.

Ricard-Blum S., and Ruggiero F. 2005. The collagen superfamily: from the extracellular matrix to the cell membrane. Pathologie Biologie 53 :430–442. 10.1016/j.patbio.2004.12.024

Relaix F., M. Bencze, M.J. Borok, A. Der Vartanian, F. Gattazzo, D. Mademtzoglou, S. Perez-Diaz, et al. 2021. Perspectives on Skeletal Muscle Stem Cells. Nature Communications 12: 692. 10.1038/s41467-020-20760-6.

Roche B., B. Arcangioli, and R. Martienssen. 2017. Transcriptional Reprogramming in Cellular Quiescence. RNA Biology 14:843–53. 10.1080/15476286.2017.1327510.

Sambasivan R., R. Yao, A. Kissenpfennig, L. van Wittenberghe, A. Paldi, B. Gayraud-Morel, H. Guenou, B. Malissen, S. Tajbakhsh, and A. Galy. 2011. Pax7-Expressing Satellite Cells Are Indispensable for Adult Skeletal Muscle Regeneration. Development 138:4333– 4333. 10.1242/dev.073601.

Schüler S.C., S. Dumontier, J. Rigaux, and C. Florian Bentzinger. 2021. Visualization of the Skeletal Muscle Stem Cell Niche in Fiber Bundles. Current Protocols 1:e263. 10.1002/cpz1.263.

Snijders T., J.P. Nederveen, B.R. McKay, S. Joanisse, L.B. Verdijk, L.J.C. van Loon, and G. Parise. 2015. Satellite Cells in Human Skeletal Muscle Plasticity. Frontiers in Physiology 6:283. 10.3389/fphys.2015.00283.

Su J., K. Gorse, F. Ramirez, and M.A. Fox. 2010. Collagen XIX Is Expressed by Interneurons and Contributes to the Formation of Hippocampal Synapses. The Journal of Comparative Neurology 518:229–53. 10.1002/cne.22228.

Symoens S., Renard M., Bonod Bidaud C., Syx D., Vaganay E., Malfait F., Ricard-Blum S., Kessler E., van Laer L., Coucke P., Ruggiero F., De Paepe A. 2010. Identification of binding partners interacting with the α1-N-propeptide of type V collagen. Biochemical Journal 433:371–81.

Theret M., F.M.V. Rossi, and O. Contreras. 2021. Evolving Roles of Muscle-Resident Fibro-Adipogenic Progenitors in Health, Regeneration, Neuromuscular Disorders, and Aging. Frontiers in Physiology 12 673404. 10.3389/fphys.2021.673404.

Tsutsui K., R-I Manabe, T. Yamada, I. Nakano, Y. Oguri, D.R. Keene, G. Sengle, L.Y. Sakai, and K.Sekiguchi. 2010. ADAMTSL-6 Is a Novel Extracellular Matrix Protein That Binds to Fibrillin-1 and Promotes Fibrillin-1 Fibril Formation. Journal of Biological Chemistry 285: 4870–82. 10.1074/jbc.M109.076919.

Tucić M., V. Stamenković, and P. Andjus. 2021. The Extracellular Matrix Glycoprotein Tenascin C and Adult Neurogenesis. Frontiers in Cell and Developmental Biology 9:674199. 10.3389/fcell.2021.674199.

Urciuolo A., M. Quarta, V. Morbidoni, F. Gattazzo, S. Molon, P. Grumati, F. Montemurro, et al. 2013. Collagen VI Regulates Satellite Cell Self-Renewal and Muscle Regeneration. Nature Communications 4:1964. 10.1038/ncomms2964.

Van Rijn A., L. Paulis, J. te Riet, A. Vasaturo, I. Reinieren-Beeren, A. van der Schaaf, A.J. Kuipers, et al. 2016. Semaphorin 7A Promotes Chemokine-Driven Dendritic Cell Migration. The Journal of Immunology 196:459–68. 10.4049/jimmunol.1403096.

Van Velthoven C.T.J., and T.A. Rando. 2019. Stem Cell Quiescence: Dynamism, Restraint, and Cellular Idling. Cell Stem Cell 24: 213–25. 10.1016/j.stem.2019.01.001.

Verma M., Asakura Y., B.S.R. Murakonda, T. Pengo, C. Latroche, B. Chazaud, L.K. McLoon, and A. Asakura. 2018. Muscle Satellite Cell Cross-Talk with a Vascular Niche Maintains Quiescence via VEGF and Notch Signaling. Cell Stem Cell 23:530–543.e9. 10.1016/j.stem.2018.09.007

Verhagen M.G., and R.J. Pasterkamp. 2020. Axon guidance: semaphorin/neuropilin/plexin signaling. In Cellular Migration and Formation of Axons and Dendrites (Second Edition). Academic Press. Chapter 5, Pages 109–122.

Webster M.T., Manor U., Lippincott-Schwartz J., Fan C.-M. 2016. Intravital Imaging Reveals Ghost Fibers as Architectural Units Guiding Myogenic Progenitors during Regeneration. Cell Stem Cell 18:243–252. 10.1016/j.stem.2015.11.005

Yang C., H. Yu, R. Chen, K. Tao, L. Jian, M. Peng, X. Li, M. Liu, and S. Liu. 2019. CXCL1 Stimulates Migration and Invasion in ER-negative Breast Cancer Cells via Activation of the ERK/MMP2/9 Signaling Axis. International Journal of Oncology 55:684–696. 10.3892/ijo.2019.4840.

